# Hyaluronan Surface Architecture Dictates Colorectal Cancer Progression and Extracellular Vesicle Communication

**DOI:** 10.64898/2026.09.16.751988

**Authors:** Debashish Paul, Neha Mathur, Tanya Agrawal, Sunita Saha, Natasha Natasha, Suchetan Pal, Anil Kumar Dasanna, Tatini Rakshit

**Affiliations:** Department of Chemistry, Shiv Nadar Institution of Eminence, Delhi-NCR, Uttar Pradesh 201314, India; Department of Physical Sciences, Indian Institute of Science Education and Research Mohali, SAS Nagar 140306, Punjab, India; Department of Chemistry, Indian Institute of Technology Bhilai, Durg, Chhattisgarh 491001, India; Department of Bioscience and Biomedical Engineering, Indian Institute of Technology Bhilai, Durg, Chhattisgarh 491001, India

## Abstract

Hyaluronan (HA) is a principal component of the tumor glycocalyx in colorectal cancer (CRC). However, how the disease progression is linked to HA abundance and its nanoscale organization remains unclear. Single-molecule measurements of surface glycans on cell membranes and extracellular vesicles (EVs) have not yet been correlated. In this work, using single-molecule force spectroscopy, we mapped HA density and chain length on CRC cells and their EVs across Dukes’ stages. HA density increased with stage in both compartments, but their organization diverged. Cell-surface HA became progressively fragmented, whereas EVs remained enriched in short HA chains at every stage. EVs, therefore, appear to select HA during formation rather than inherit it from the parent cell. This divergence had mechanical consequences. Both cells and EVs softened with stage, and removing HA reversed this softening. In addition, coarse-grained membrane simulations revealed that both HA chain length and surface density regulate membrane wrapping, with chain length primarily influencing wrapping kinetics and surface density affecting the final wrapping extent. These findings provide a physical basis for the differences we observed in EV uptake. Reprogramming stage D cells with exogenous high-molecular-weight HA reversed this signature, lowering EV surface HA density, stiffening the vesicles, slowing migration, and suppressing EV uptake by recipient cells. These findings establish HA surface architecture as a stage-encoded and experimentally reversible determinant of CRC progression.

---

Keywords: Colorectal cancer, extracellular vesicles, hyaluronan, surface density, contour length, viscosity, elasticity, atomic force microscopy

## Introduction

Hyaluronan (HA) is a ubiquitous, high-molecular-mass polysaccharide of the vertebrate extracellular matrix (ECM) that governs tissue hydration, viscoelasticity, and macromolecular organization through its expanded, semi-flexible random-coil conformation^1^. Beyond passive space-filling, HA signals through cell-surface receptors, including CD44 and RHAMM, assembling into a receptor-anchored pericellular coat that mediates mechanotransduction and cell-ECM communication^2^. This homeostasis reflects a dynamic balance between synthesis by hyaluronan synthases (HAS1–3) and degradation by hyaluronidases^3^, a balance that is characteristically disrupted in malignancy.

Colorectal cancer (CRC) is the third most commonly diagnosed malignancy worldwide, with incidence projected to rise nearly 60% by 2030 and prognosis remaining strongly stage-dependent^4^. Identifying stage-specific molecular and biophysical alterations within the tumor microenvironment is therefore central to improving early detection and prognostic stratification^5,6^. CRC tissue consistently shows elevated HA relative to normal colonic epithelium, but accumulating evidence indicates that progression is defined less by HA abundance than by systematic remodeling of its molecular architecture^7,8^. In early disease, upregulated HAS2/HAS3 activity increases high-molecular-weight HA (HMW-HA) at the cell surface, sustaining CD44-driven proliferation and motility, while concurrent hyaluronidase activity generates low-molecular-weight HA (LMW-HA) fragments that promote angiogenesis and inflammation^9–11^. As the disease advances, overexpression of Hyal-1/2/3 shifts this balance further, depleting long-chain HA and enriching circulating LMW-HA fragments that correlate with lymphovascular invasion, nodal spread, and relapse^12^. Total HA thus remains elevated throughout progression, but its chain-length architecture is progressively reorganized, a distinction with direct mechanical consequences, since HA’s hydrated polymer architecture is a primary determinant of glycocalyx stiffness, steric interactions, and cell deformability. Whether this architectural remodeling, as distinct from abundance, directly governs the nanomechanical phenotype of CRC cells has not been tested.

CRC cells additionally communicate through extracellular vesicles (EVs), nanoscale, lipid-bilayer particles that transfer proteins, nucleic acids, and surface carbohydrates between cells and are increasingly pursued as CRC biomarkers^13,14^. Tumor-derived EVs frequently carry HA-rich coats implicated in immune evasion, matrix remodeling, and pre-metastatic niche formation^15^, and HA’s interactions with CD44, RHAMM, and endosomal trafficking machinery position it as a plausible regulator of EV biogenesis and cargo selection^16,17^. Yet quantitative, single-molecule-resolution characterization of HA, indeed, of surface glycans and receptors generally, has been confined mostly to cell membranes^18,19^, with a few reports on EVs^20,21^. Importantly, whether EV-surface HA architecture tracks, diverges from, or is mechanistically decoupled from the HA remodeling occurring at the parent cell surface remains entirely unknown.

This is a consequential gap rather than a technical afterthought. If EV-surface HA architecture simply mirrors the parent cell, an inert carryover of membrane composition during budding, then HAs would be expected to fragment in step with the cell, and any diagnostic or mechanistic value of EV-surface HA would be redundant with cell-surface measurements already established in the literature. If, instead, EV biogenesis actively selects for or otherwise reorganizes HA independently of the parent cell’s chain-length state, EV-surface HA would constitute a distinct, non-redundant axis of tumor biology with direct consequences for how EVs engage recipient-cell membranes, since HA density and chain length are themselves physical determinants of membrane wrapping and internalization kinetics. Resolving which of these scenarios holds requires simultaneous, quantitative, single-molecule-resolution mapping of HA density and chain length on cells and their EVs across defined stages of disease.

Here, we address this gap by integrating AFM-based single-molecule force spectroscopy, nanomechanical depiction, coarse-grained membrane simulation, and functional cellular assays across CRC lines spanning Duke stage B (SW480, HT29) and stage D (Colo-205, HCT116), anchored by two specificity controls: a patient-matched stage B-to-stage C isogenic pair (SW480/SW620), which isolates stage-dependent remodeling from inter-patient genetic variability, and EVs from non-malignant colonic fibroblasts (CCD-18Co), which establish that the phenotype is cancer-associated rather than a general EV property. We show that HA surface density rises steeply with stage on both cells (0.56-10.6%) and EVs (3.7-17.9%) but that the two compartments diverge in architecture: cell-surface HA chain length contracts progressively with stage, from micron-scale to sub-200 nm, while EV-associated HA remains low-molecular-weight-enriched and structurally complex regardless of stage, indicating that EV biogenesis imposes its own, parent-cell-independent selection on HA chain length rather than passively inheriting it. We further show that this architecture is mechanically contributory not incidental: enzymatic HA removal or synthesis inhibition increases cell stiffness by nearly two orders of magnitude and EV stiffness by roughly twofold, directly linking HA organization to the biomechanical softening long associated with malignant progression^22^. To ask whether this architecture has direct physical consequences for EV-cell engagement, we used coarse-grained molecular dynamics simulations of polymer-grafted vesicles interacting with a fluid membrane. The simulations show that HA density and chain length jointly determine wrapping efficiency, providing a physical basis for the differential EV uptake we observe experimentally. Finally, we show that this entire signature is reversible: supplementing stage D cells with exogenous high-molecular-weight HA lowers EV surface density, stiffens EVs, slows migration, and suppresses EV uptake by recipient cells, thereby reprogramming a stage D cell toward a stage B-like phenotype across all measured axes. Together, these results identify HA surface architecture at the cell-EV interface as a stage-encoded, mechanically contributory, and therapeutically reversible driver of CRC progression, and nominate EV-surface HA architecture as a biomarker and engineerable target distinct from HA abundance alone. More broadly, these findings suggest that the pericellular and extracellular vesicle glycocalyx should be evaluated as an architecturally dynamic, mechanically active compartment of the tumor microenvironment, rather than a passive structural reservoir of a single overexpressed biomolecule.

## Results

### Isolation and characterization of EVs

EVs were isolated from CRC cell lines spanning distinct disease stages by differential ultracentrifugation (**Fig. S1a**) and validated by three complementary techniques: TEM confirmed uniformly distributed, near-spherical vesicles (**Fig. S1b**); NTA revealed a predominantly small-EV-range particle population and showed stage-dependent variation in both concentration and mean diameter (**Fig. S1c; Table S1**). Bead-assisted flow cytometry confirmed expression of the canonical markers CD9, CD63, and CD81 (**Fig. S1d**)^23^. These vesicles, therefore, constitute a validated, stage-resolved platform for the subsequent HA characterization.

### HA is elevated on both cells and EVs with CRC progression, but ensemble methods cannot resolve its architecture

Confocal microscopy showed predominantly membrane-localized HA that increased progressively from stage B to stage D CRC cells (**Fig. 1A, B**), corroborated by FACS on both cells and their corresponding EVs (**Fig. 1C**). These ensemble measurements, however, lack the sensitivity and spatial resolution to quantify HA at the single-molecule level. We therefore used single-molecule force spectroscopy (SMFS) to directly resolve the HA architecture.

**Figure 1.**
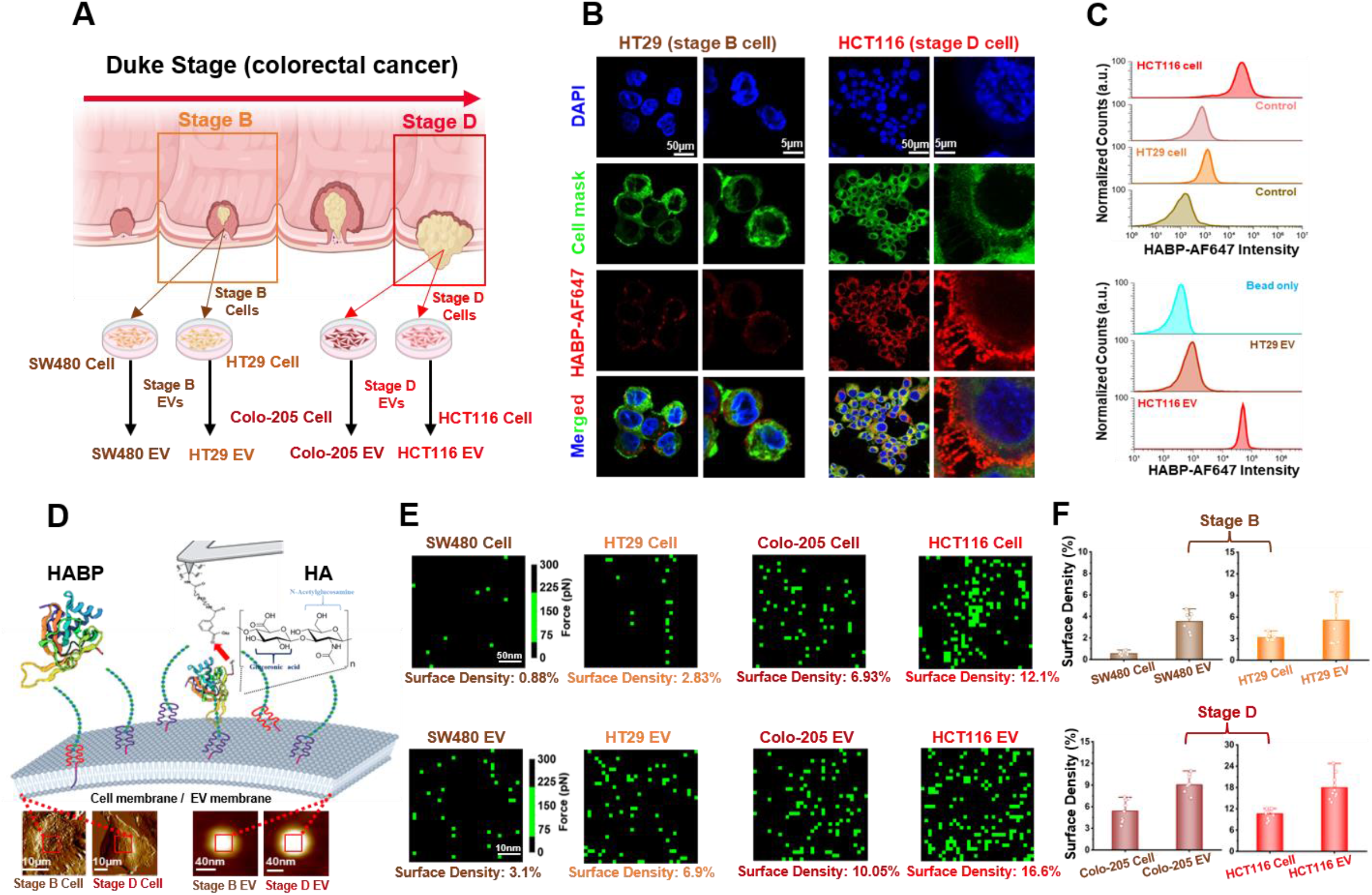
Stage-dependent HA expression on CRC cells, EVs, and quantitative nanoscale mapping of HA surface density. (**A**) Schematic illustration of the experimental design. Human CRC cell lines representing Dukes’ stage B (SW480 and HT29) and Dukes’ stage D (Colo-205 and HCT116) were cultured, and EVs were isolated from conditioned media by differential ultracentrifugation. (**B**) Representative confocal microscopy images showing HA expression on CRC cells (stage B and D) following staining with HABP-AF647 (red). Cell nuclei were counterstained with DAPI (blue), and the plasma membrane was labeled with CellMask™ (green). Merged images demonstrate increased HA abundance in the stage D HCT116 cells compared with the stage B HT29 cells. (**C**) Flow cytometry histograms validating specific binding of HABP-AF647. Upper panels compare fluorescence intensity distributions of HT29 and HCT116 cells with their respective unstained/negative controls. Lower panels show fluorescence intensity of isolated HT29 and HCT116 EVs relative to bead-only controls. Rightward shifts in fluorescence intensity confirm specific HA detection and indicate increased HA expression in stage D-derived cells and EVs. (**D**) Schematic representation of AFM-based SMFS for quantitative measurement of cell/EV surface HA density. AFM cantilevers were functionalized with HABP through flexible PEG linkers, enabling specific interaction with HA molecules present on cell and EV membranes. Insets show representative AFM topographic images of stage B and stage D cells and individual EVs selected for SMFS measurements. (**E**) Representative adhesion-force density maps obtained from AFM-SMFS measurements of individual CRC cells (top row) and their corresponding EVs (bottom row). Green pixels represent specific HABP–HA binding events detected during force-volume mapping. Surface HA density progressively increased from SW480 to HT29, Colo-205, and HCT116 in both cells and EVs. Representative quantified HA surface densities are indicated below each map. (**F**) Quantitative comparison of HA surface density measured by AFM-SMFS for CRC cells and their corresponding EVs, across different stages (upper panel, stage B and lower panel, stage D). Both parental cells and EVs exhibit a stage-dependent increase in HA surface density, with the highest levels observed in stage D-derived HCT116 samples. Data are presented as mean ± SD from 10 analyzed cells/EVs.

### Single-molecule AFM reveals a stage-graded increase in HA surface density on cells and EVs, independent of patient origin

FD-based AFM was performed using HABP-functionalized probes whose specificity was independently validated by SMFS^20^ (**Fig. S2c**). The HA–HABP interaction followed the Bell-Evans model (k_off_ = 0.008 s⁻¹, reactive compliance x_β_ = 0.13 nm; **Fig. S2a–b**), and specific single-molecule adhesion events (35–200 pN) were detected on every CRC cell line and EV type but absent on BSA-coated controls, confirming specificity (**Fig. S2c**). Force-volume adhesion mapping (**Fig. 1D**) showed HA distributed heterogeneously across both surfaces, with localized nanoscale clustering (**Fig. 1E**); we therefore quantified HA expression as the percentage of HA-specific adhesion events per mapped area (surface density).

Quantitative force maps revealed a clear, monotonic stage-dependent increase in HA surface density on both cells and EVs (**Fig. 1F; Table S1**). Stage B cells showed sparse coverage 0.56 ± 0.10% (SW480) and 3.17 ± 0.40% (HT-29), rising to 6.7 ± 2.7% (Colo-205) and 10.6 ± 1.4% (HCT116) at stage D. EVs followed the same trend at consistently higher magnitude: 3.7 ± 0.8% (SW480 EV) and 5.5 ± 2.8% (HT-29 EV) at stage B, rising to 9.3 ± 4.8% (Colo-205 EV) and 17.9 ± 3.3% (HCT116 EV) at stage D. EVs from the non-malignant CCD-18Co line showed the lowest density of any sample (0.4 ± 0.1%), consistent with flow cytometry (**Fig. S9a–c**) and establishing that this is a cancer-associated rather than general EV property.

Since cell and EV surface areas differ by orders of magnitude, we report density as the normalized metric of comparison throughout. The corresponding estimated absolute HA molecule counts (**Table S1**) show that, despite carrying far fewer total HA molecules, EVs consistently present a higher “local” HA density than their parent cells. Expression of CD44, the principal HA receptor, increased with stage on cells but showed no significant stage-dependent difference on EVs (**Fig. S10**). Measured HA–HABP rupture forces agreed with prior reports, further supporting measurement reliability^20,21^.

To rule out inter-patient variability as a confound, we performed the identical measurement on SW620 cells-a stage C metastatic line derived from the same patient as stage B SW480. SW620 density (3.9 ± 0.70%) was significantly higher than that of SW480 (0.56 ± 0.10%; **Fig. S6a–b**), confirming that the increase in density tracks tumor progression rather than genetic background.

### HA surface density is mechanistically linked to CRC progression, not merely correlated with it

To test whether FD-AFM density values reflect a mechanistically modifiable property rather than a fixed cell-type signature, we enzymatically removed HA (hyaluronidase) or inhibited its biosynthesis (4-methylumbelliferone, 4-mU) in HCT116 cells, the line with the highest baseline HA density, and re-isolated EVs from the treated cells (**Fig. S5a, a**). Both treatments preserved EV integrity, morphology, and size distribution (**Fig. S5a, f–g**) while producing large, concordant reductions in HA density by confocal, FACS, and FD-AFM (**Fig. S5a, b–e, h–j**). On cells, density dropped from 10.6 ± 1.4% (untreated) to 3.0 ± 2.7% (4-mU) and 1.15 ± 0.6% (hyaluronidase); on EVs, from 17.9 ± 3.3% to 9.2 ± 2.8% and 2.6 ± 0.6%, respectively. This bidirectional, dose-graded response across two independent perturbation mechanisms confirms that FD-AFM density is a specific, sensitive, and engineerable readout of surface HA, a prerequisite for the subsequent mechanical and functional experiments.

### Importantly, the HA chain length shortens progressively with cell stage, but EVs retain low-molecular-weight HA regardless of parental stage

SMFS contour-length (L_c_) analysis of HA–HABP unbinding events, fit to the worm-like-chain model (persistence length 4.1–4.4 nm, consistent with reported single-HA-molecule values^24^), showed a clear architectural transition (**Fig. 2A**). Stage B cells carried predominantly long HA chains, Gaussian maxima at 0.49, 1.25, and 2.09 µm (SW480) and 0.88, 1.72, and 2.87 µm (HT-29), while stage D cells shifted sharply toward short chains: 0.12 and 0.31 µm (Colo-205) and 0.09, 0.19, and 0.35 µm (HCT116) (**Fig. 2B**). Consistent with this, single-rupture events (indicating isolated chains) dominated stage B force curves, while multi-rupture events (indicating denser, more clustered chains) became markedly more frequent at stage D (**Fig. S3a–f**).

**Figure 2.**
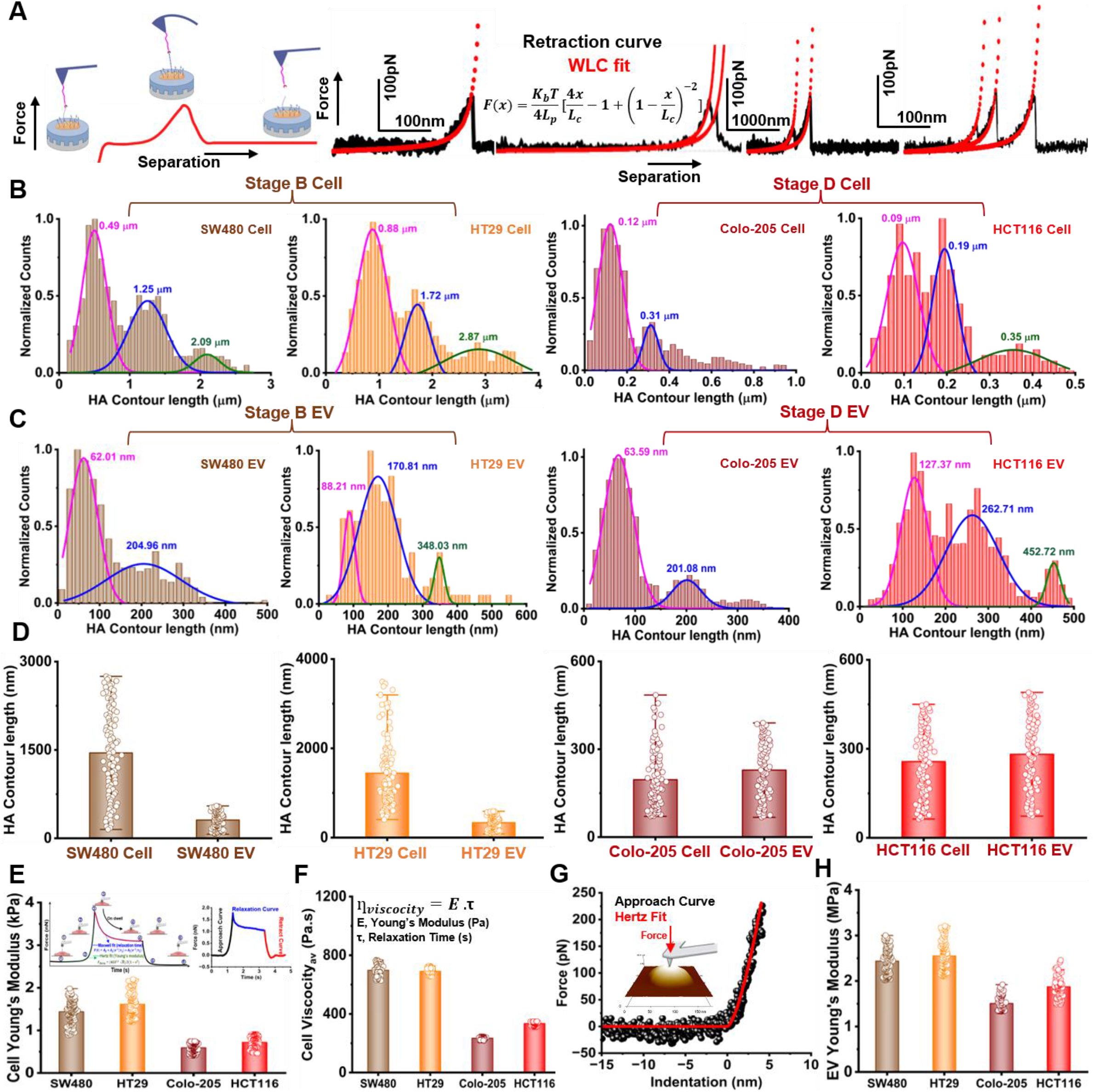
Stage-dependent remodeling of HA contour lengths and elasticity of CRC cells and EVs. (**A**) Schematic representation of AFM-based SMFS used to measure HA contour length on the surface of CRC cells and EVs. During cantilever retraction, individual HA chains bind to the HABP-functionalized AFM probe and are sequentially stretched, producing characteristic rupture events in the force–distance (retraction) curves. Representative force-extension curves obtained from stage B and stage D cells and EVs are shown. The retract curves were fitted using the worm-like chain (WLC) model (red), to extract HA contour length (Lc) for individual molecular interactions. (**B**) Probability distributions of HA contour lengths measured on the surface of CRC cells. Histograms were constructed from WLC-fitted contour lengths and deconvoluted into multiple Gaussian populations representing distinct HA chain-length subpopulations. Stage B cells (SW480 and HT29) exhibited a substantial population of long HA chains extending to approximately 1–3 μm, whereas stage D cells (Colo-205 and HCT116) were dominated by significantly shorter HA chains, with contour lengths below 0.5 μm. (**C**) Probability distributions of HA contour lengths measured on isolated CRC EVs. EV-associated HA exhibited contour-length distributions around 60-450 nm. (**D**) Comparison of HA contour length between cells and EVs across different CRC stages (from Stage B to Stage D, left to right). (E) Schematic illustration of AFM nanoindentation used to quantify the viscoelastic behavior. Force–time curves were acquired during probe approach, dwell period, and retraction. During the constant-force dwell period, time-dependent deformation was analysed using a standard viscoelastic model to determine the Young’s modulus (E) and relaxation time (τ). Representative fitting equations used for the viscoelastic analysis are shown (**inset**). Representative force–time curve obtained during AFM nanoindentation illustrating the approach (black), dwell (blue), and retraction (red) segments used for viscoelastic fitting and extraction of relaxation parameters (**inset**). Quantification of the apparent Young’s modulus of CRC cells measured by AFM nanoindentation (**E**). Quantification of cell viscosity was calculated from the relationship η = E × τ, where *E* is the Young’s modulus and *τ* is the relaxation time (**F**). Representative AFM force–indentation curve acquired during nanoindentation of EVs. The approach curve was fitted using the Hertz contact model (red) (**G**), Young’s modulus of CRC EVs (**H**).

EV-associated HA did not follow this trajectory. Regardless of parental stage, EVs displayed consistently short contour lengths in the range of tens to a few hundred nanometers: 62.0 and 205.0 nm (SW480 EV), 88.2, 170.8, and 348.0 nm (HT-29 EV), 63.6 and 201.1 nm (Colo-205 EV), and 127.4, 262.7, and 452.7 nm (HCT116 EV) (**Fig. 2C**). Even stage B-derived EVs therefore carry an HA architecture that resembles stage D cells far more than it resembles their own parent cell surface (**Fig. 2D**). Quantifying the HMW: LMW-HA ratio from these distributions (**Table S1**) sharpens this picture: stage B cells are HMW-HA-dominant, stage D cells are almost completely LMW-HA-dominant, but EVs at “both” stages are predominantly LMW-HA-enriched. The same conclusion held in the isogenic pair: SW620 chains (Gaussian maxima 0.21, 0.76, 1.38 µm) were significantly shorter than SW480 (0.49, 1.25, 2.09 µm), and the HMW: LMW ratio shifted from 86:14 to 50:50 (**Fig. S6c–d; Table S1**), again disconnecting this remodeling from patient background.

Because LMW-HA fragments function as damage-associated molecular patterns capable of driving inflammatory and pro-metastatic signaling^25^, the consistent LMW-HA enrichment on EVs independent of parental chain-length state suggests that EV biogenesis actively selects for, rather than passively inherits, short HA chains.

### Coarse-grained simulations link HA chain length and density to vesicle–membrane wrapping efficiency and membrane bending energy

To test whether HA surface architecture has direct physical consequences for EV–cell engagement, we modelled a rigid spherical vesicle decorated with grafted semiflexible polymers, with grafted chain length representing HA molecular weight and grafting density corresponding to the FD-AFM-measured surface density. This modelling framework builds on previous theoretical and computational studies of membrane wrapping, which established the importance of the interplay between particle–membrane adhesion, membrane deformation, and the properties of polymers grafted to particle surfaces^26–30^. The use of semiflexible grafted polymers is also consistent with recent coarse-grained models demonstrating that filamentous structures on particle and cell surfaces can strongly influence membrane wrapping dynamics^31,32^. The vesicle was placed in contact with a one-particle-thick, solvent-free fluid membrane (**Fig. 3A**). We quantified membrane wrapping W(t) as the fraction of grafted polymer monomers adhered to membrane particles:

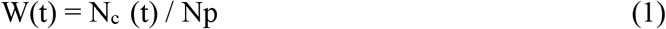

where Np is the total number of grafted polymer monomers and N_c_ (t) is the number of polymer monomers adhered to membrane particles at time t. Representative trajectories at φs = 0.20 (**Fig. 3B**) showed initial contact, beginning of wrapping, half wrapping, and complete wrapping. Short-chain vesicles reached high wrapping fractions more rapidly, whereas longer chains wrapped more slowly and frequently remained partially wrapped. At fixed chain length (L = 20σ), wrapping saturation decreased with increasing grafting density: sparse coverage (φs = 0.05) produced near-complete wrapping (W ≈ 0.9), whereas dense coverage (φs = 0.20) resulted in a lower saturation value (W ≈ 0.75) (**Fig. 3C**), consistent with increased steric resistance from closely packed chains. At fixed grafting density (φs = 0.20), wrapping decreased with increasing chain length: short chains (L = 10σ) reached saturation rapidly (W ≈ 0.77), whereas long chains (L = 40σ) wrapped more slowly and remained incompletely wrapped at the end of the simulations (**Fig. 3D**). The combined (L, φs) phase map (**Fig. 3E**) shows that time-averaged wrapping efficiency ⟨W⟩ is maximal (≈0.9) for short, sparse chains and falls below 0.5 for long, dense chains, with chain length the more sensitive of the two parameters.

**Figure 3.**
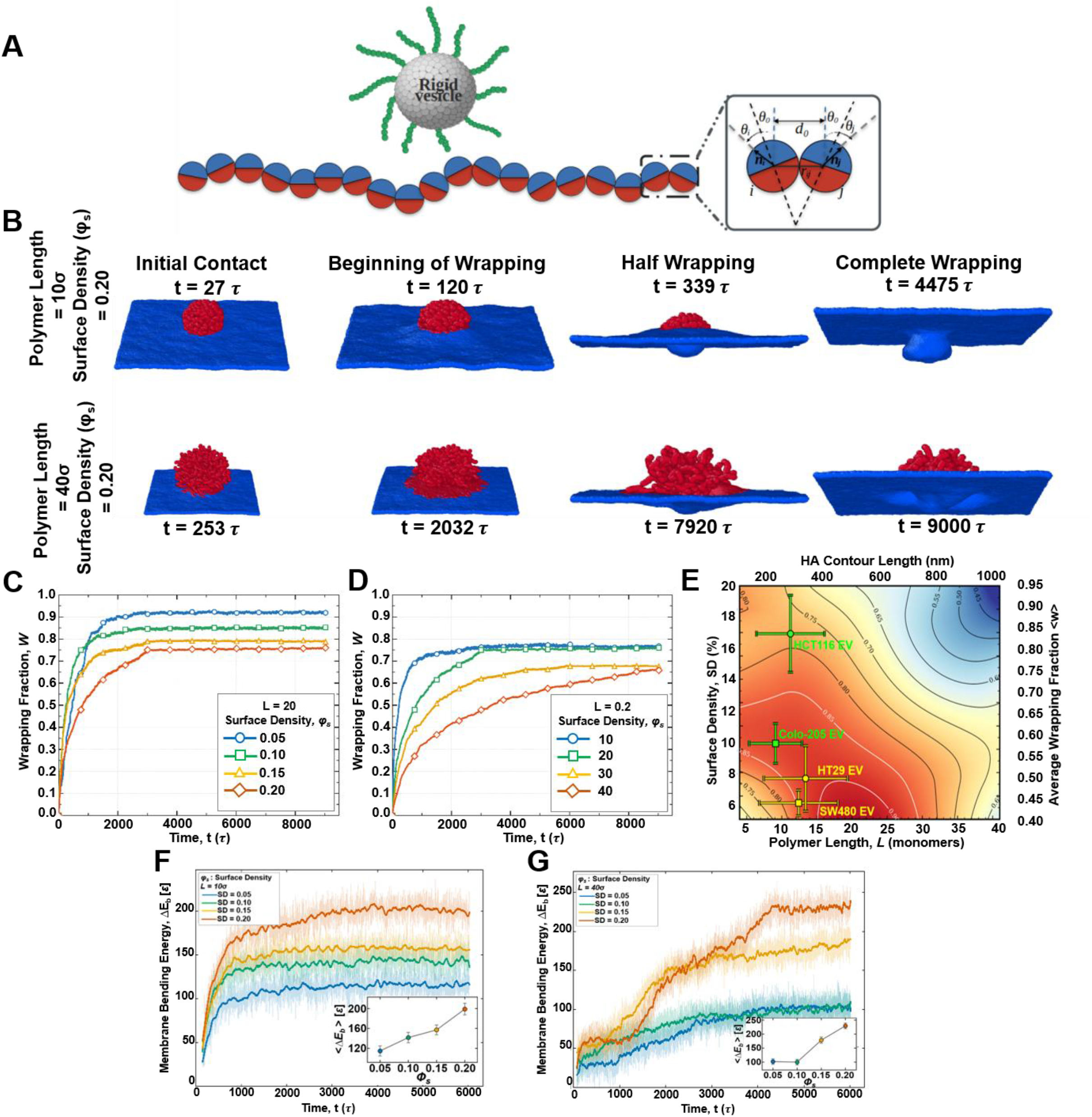
Coarse-grained simulations reveal that HA surface density and chain length dictate membrane wrapping dynamics and bending energetics. **(A)** Coarse-grained model of a polymer-grafted rigid vesicle adhering to a one-particle-thick fluid membrane; the inset defines the geometry of the anisotropic membrane interaction. **(B)** Snapshots at ϕ_s_ = 0.20 for a short chain (L = 10σ, top) and a long chain (L = 40σ, bottom), shown at four stages - initial contact, beginning of wrapping, half wrapping, and complete wrapping, with the corresponding simulation times. **(C, D)** Wrapping fraction W(t) for fixed L = 20σ (a) and fixed ϕ_s_ = 0.20 (b). **(E)** Time-averaged wrapping fraction ⟨W⟩ over the (L,ϕ_s_) plane (EV systems according to their average surface density and contour length are overlaid). (F, G) Excess membrane bending energy ΔE_b_ versus time for L = 10σ **(F)** and L =40σ **(G)**.

The membrane bending energy was calculated from the dipole orientations of the membrane particles. Each membrane particle carries a dipole vector, μᵢ, oriented along the local membrane normal. Membrane curvature therefore produces an orientational mismatch between the dipoles of neighbouring particles. We quantified local membrane bending by accumulating this orientational mismatch over all neighbouring membrane-particle pairs within the deformed region surrounding the vesicle and subtracting the corresponding value obtained from an unperturbed, vesicle-free membrane simulated under identical conditions. The resulting excess bending energy, ΔE_b_, is expressed in units of the membrane interaction energy, ε, and represents the additional curvature energy stored in the membrane as a consequence of the wrapping event. For short chains (L = 10σ; **Fig. 3F**), ΔE_b_ increased steeply and reached a well-defined plateau, whereas for long chains (L = 40σ; **Fig. 3G**), it increased much more gradually and reached saturation only at late times, consistent with the slower and incomplete wrapping of long-chain vesicles. In both cases, the plateau value increased monotonically with grafting density (insets, **Fig. 3F–G**), indicating that a denser grafted layer requires the membrane to adopt a more strongly curved conformation to establish contact with the vesicle. At a fixed grafting density, however, the plateau bending energy was consistently lower for L = 40σ than for L = 10σ, because the thicker grafted layer sterically opposes membrane engulfment, preventing the membrane from reaching the highly curved, near-enclosing geometry attained in the short-chain systems. Diffusion analysis in the absence of a membrane (**Fig. S4a-b**) showed that the calculated diffusion coefficients decreased with increasing chain length and grafting density, indicating greater mobility for shorter and more sparsely grafted vesicles (see Supporting Information for the MSD analysis).

Together, these simulations show that HA chain length, more than chain density, controls how efficiently a vesicle wraps against a membrane, providing a physical basis for HA-dependent EV uptake.

### Cell and EV nanomechanics soften progressively with CRC stage and are directly governed by HA

AFM topographical analysis of height, diameter, roughness, and surface area (**Table S1**) revealed stage-dependent morphological changes in both cells and EVs, with the latter retaining spherical morphology throughout. Using an AFM dwell-retract stress-relaxation protocol (**Fig. 2E**; Hertz fit for Young’s modulus, second-order Maxwell fit for relaxation times τ1/τ2), stage B cells relaxed more slowly than stage D cells: τ_1_ = 0.08 ± 0.02 s / τ_2_ = 0.81 ± 0.1 s (HT-29) and τ_1_ = 0.08 ± 0.02 s / τ_2_ = 0.92 ± 0.2 s (SW480), versus τ_1_ = 0.072 ± 0.01 s / τ_2_ = 0.85 ± 0.2 s (HCT116) and τ_1_ = 0.07 ± 0.01 s / τ_2_ = 0.74 ± 0.1 s (Colo-205) (**Fig. S7a–d**). Average relaxation time dropped accordingly with stage (0.55–0.50 s at stage B to 0.40–0.45 s at stage D; **Fig. S7e**).

Young’s modulus dropped by roughly 50% from stage B to stage D, 1.6 ± 0.3 kPa (HT-29) and 1.4 ± 0.2 kPa (SW480) versus 0.72 ± 0.17 kPa (HCT116) and 0.59 ± 0.1 kPa (Colo-205) (**Fig. 2E**) and, since viscosity scales with the product of modulus and relaxation time, average viscosity also dropped from 698 ± 41 Pa·s (SW480) / 691 ± 18 Pa·s (HT-29) to 334 ± 12 Pa·s (HCT116) / 235 ± 8 Pa·s (Colo-205), with both the fast and slow viscoelastic components (η1, η2) following the same trend (**Fig. 2F; Fig. S7f-g)**. EV stiffness showed the identical directional trend at a smaller magnitude: 2.6 ± 0.3 MPa (HT-29 EV) / 2.4 ± 0.3 MPa (SW480 EV) at stage B versus 1.9 ± 0.3 MPa (HCT116 EV) / 1.6 ± 0.2 MPa (Colo-205 EV) at stage D (**Fig. 2G-H**).

Critically, this softening was not merely coincident with HA remodeling; it was reversible by direct HA manipulation. Enzymatic or pharmacological HA removal significantly increased both cellular viscosity (**Fig. S5b, a–c**) and EV stiffness (**Fig. S5b, a, b, d**), and the isogenic SW480–SW620 pair reproduced the same relationship independent of genetic background (SW620 EVs significantly softer than SW480 EVs; **Fig. S6e**). HA surface density and chain length are therefore direct, contributing determinants of CRC cell and EV nanomechanics, not passively correlated with stage.

### Stage-dependent HA remodeling is conserved on microvesicles

To test whether this remodeling is specific to exosome-enriched EVs or a general feature of secreted vesicles, we analyzed microvesicles (MVs), which bud directly from the plasma membrane and inherit its composition more directly (**Fig. S8b, a-b**). Stage D (HCT116) MVs showed higher and more uniform HA density (6.1 ± 1.7%) than stage B (HT-29) MVs (4.5 ± 0.8%; **Fig. S8b, d–f**), abolished by 4-mU treatment (0.5 ± 0.1%; **Fig. S8b, c**), confirming specificity. Contour-length analysis showed the same architectural shift as small EVs: HT-29 MVs carried two populations (147.9, 334.6 nm) while HCT116 MVs carried four, all shorter on average (59.3, 127.2, 261.3, 403.0 nm; **Fig. S8b, e, g**), with an LMW: HMW ratio of ∼100:0 for both. Stage-dependent HA remodeling toward higher density and shorter chains is therefore conserved across vesicle biogenesis routes.

### Exogenous HMW-HA reprograms stage D cell migration and EV architecture

Given the HMW-HA dominance of stage B cells versus the LMW-HA dominance of stage D cells, we tested whether exogenously supplementing stage D (HCT116) cells with FITC-labelled HMW-HA (∼1500 kDa) could remodel their glycocalyx and reverse their phenotype. After 24 - 48 h incubation, confocal imaging showed colocalization of exogenous HMW-HA with native surface HA (**Fig. 4A–B; Fig. S11**), and flow cytometry showed a progressive decline in endogenous LMW-HA alongside a rise in surface HMW-HA and pericellular coat thickness (**Fig. 4C–D; Table S1**), indicating partial replacement rather than simple addition.

**Figure 4.**
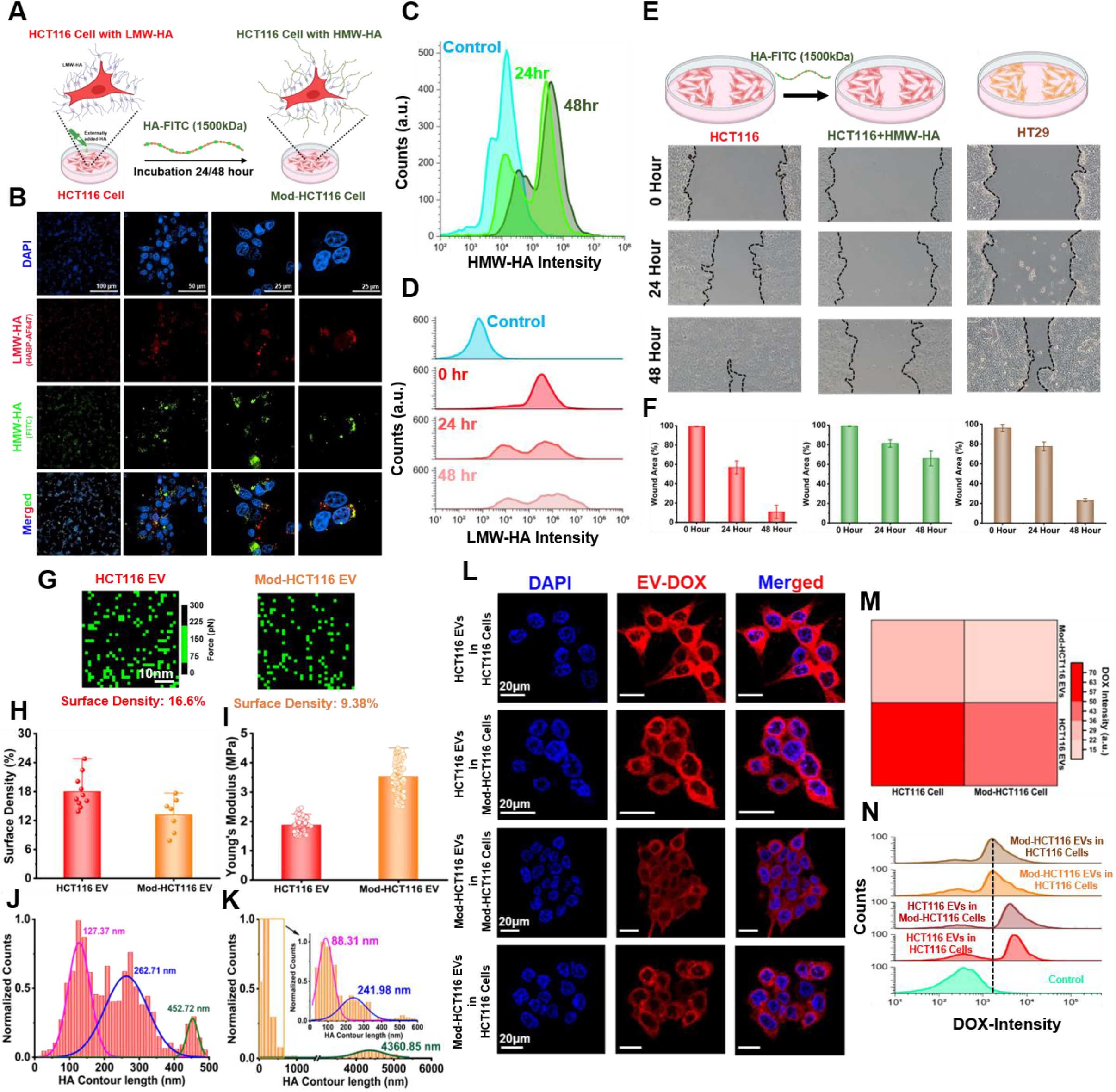
Replacement of LMW-HA with HMW-HA reprogrammes the HCT116 CRC migration, remodels EV surface biophysical properties, and attenuates EV uptake. **(A)** Schematic illustration of the experimental workflow for engineering HCT116 cells by HMW-HA (HA-FITC, 1500 kDa). The cells were incubated with HA-FITC for 24 or 48 h to generate HMW-HA-modified HCT116 (Mod-HCT116) cells, while untreated HCT116 cells served as controls. **(B)** Confocal microscopy images of HCT116 and Mod-HCT116 cells showing nuclei stained with DAPI (blue), endogenous low-molecular-weight hyaluronan (LMW-HA; red), exogenous HMW-HA-FITC (green) **(C)** Flow cytometry of HMW-HA-FITC treated HCT-116 cells, progressive rightward shift in fluorescence intensity after 24 and 48 h relative to untreated control cells with increased surface accumulation of HMW-HA over time was noticed (**D**) Flow cytometry of endogenous LMW-HA expression on HCT116 cells before (0 h) and after 24 and 48 h of HMW-HA treatment, a gradual decrease in LMW-HA fluorescence intensity was observed. (**E**) Representative bright-field images from scratch wound-healing assays comparing migration of untreated HCT116 cells, Mod-HCT116 cells, and HT29 cells at 0, 24, and 48 h following wound generation. Mod-HCT116 cells exhibited delayed wound closure relative to untreated HCT116 cells, whereas HT29 cells displayed an intermediate migration phenotype. Dashed lines indicate wound boundaries. (**F**) Quantitative analysis of wound closure area (%) for HCT116, Mod-HCT116, and HT29 cells at different time points (**G**) Representative AFM-based adhesion maps of EVs isolated from HCT116 and Mod-HCT116 EVs, showing spatial distribution of HA-unbinding events on individual EV surfaces. Surface HA density (%) calculated from the adhesion maps is indicated below each image. (**H**) Quantification of EV surface HA density determined from AFM single-molecule force spectroscopy measurements (**I**) Young’s modulus of HCT116 EVs and Mod-HCT116 EVs obtained from AFM nanoindentation measurements (**J**) Contour length distribution of HA molecules measured on native HCT116 EVs obtained by fitting AFM force-extension curves using the WLC model, revealing distinct HA populations with contour lengths 127-453 nm. (**K**) Contour length distribution of HA on Mod-HCT116 EVs showing contour length populations of 88 -242 nm (inset). A small fraction of events exhibited contour lengths of approximately 4300 nm, consistent with the presence of exogenously introduced HMW-HA on the EV surface. (**L**) Confocal microscopy images showing uptake of doxorubicin (DOX)-loaded EVs by HCT116 cells and Mod-HCT116 cells. Cells were incubated with either native HCT116 EVs or Mod-HCT116 EVs loaded with DOX. Nuclei were stained with DAPI (blue), DOX fluorescence is shown in red. (M) Heatmap showing the quantification of intracellular DOX intensity following EV uptake. (N) FACS of DOX intensity showing reduced cellular uptake of Mod-HCT116 EVs relative to native HCT116 EVs.

Functionally, untreated stage D cells showed the highest migratory capacity in scratch assays; HA degradation or synthesis inhibition reduced migration, but HMW-HA supplementation produced the largest reduction, suppressing migration to levels at or below stage B cells, with the effect strengthening over treatment duration (**Fig. 4E-F; Fig. S12**).

EVs secreted from HMW-HA-treated ("Mod-HCT116") cells were isolated and validated as intact (CD9/CD63/CD81-positive, 60–200 nm, near-spherical; **Fig. S13**). Relative to untreated stage D EVs, Mod-HCT116 EVs showed reduced HA density (13.1 ± 3.0% vs. 17.9 ± 3.3%; **Fig. 4G–H**) and increased stiffness (3.5 ± 0.5 MPa vs. 1.9 ± 0.3 MPa; **Fig. 4I**), directly reproducing, via an orthogonal perturbation, the density-stiffness relationship established by enzymatic HA removal. The chain-length distribution, however, was only partially reprogrammed: Mod-HCT116 EVs retained dominant LMW populations (88.3, 242.0 nm) but **acquired a minor, previously absent long-chain population (∼4361 nm) reflecting incorporated exogenous HMW-HA** (**Fig. 4J–K**), indicating that exogenous HMW-HA is only partially transferred onto the EV surface, while the native LMW-dominant architecture is largely retained.

This reprogramming was vesicle-type-selective. Surprisingly, MVs from HMW-HA-treated cells (**Fig. S14a-d, f**) showed no significant change in overall HA abundance (**Fig. S14e**), only modest density change (5.8 ± 1.2% vs. 6.1 ± 1.7%; **Fig. S14g–h**), and an essentially unchanged Lc distribution (105.9, 177.1 nm vs. untreated; **Fig. S14i-k**), in contrast to the significant remodeling seen in small EVs under the identical treatment.

### Modulating surface HA on either the EV or the recipient cell attenuates EV uptake

Finally, we asked whether this HA architecture regulates EV–cell communication itself. Confocal imaging confirmed EV internalization under all tested combinations of untreated/HMW-HA-modified donor EVs and untreated/HMW-HA-modified recipient cells (**Fig. 4L**). Untreated stage D EVs taken up by untreated recipient cells showed the highest uptake of any combination; modifying recipient cells with HMW-HA significantly reduced uptake of these same EVs, and EVs derived from Mod-HCT116 cells showed reduced uptake regardless of recipient cell phenotype (**Fig. 4L–N**, quantified by flow cytometry and fluorescence intensity). Modulating HA architecture at either side of the EV-cell interface donor vesicle or recipient membrane-is therefore sufficient to suppress internalization, **identifying HA surface architecture as a functional determinant of EV-mediated intercellular communication in advanced CRC.**

## Discussion

This study set out to test whether the well-established rise in tumor HA during CRC progression is better explained by abundance or by architecture, and whether that architecture is causally consequential rather than incidental. Three lines of evidence converge on the latter. First, enzymatic HA removal and biosynthesis inhibition each produced large, bidirectional, dose-graded changes in cell and EV stiffness, a level of causal control that correlation alone cannot establish. Second, the isogenic SW480-SW620 pair reproduced the density, chain length, and mechanical phenotype in the absence of any genetic differences, isolating stage from genetic background. Third, exogenous HMW-HA supplementation, an orthogonal, additive perturbation rather than a subtractive one, independently reproduced the same density–stiffness relationship and suppressed migration and EV uptake, indicating that the stage D phenotype is not a fixed cell state but an actively reversible one. Taken together, HA architecture functions as a contributing variable, allowing CRC cells and their EVs to be pushed in either direction, rather than merely serving as a passive biomarker of stage.

### Cells and EVs remodel HA independently, and this divergence is the study’s central finding

Cell-surface HA contour length contracted by roughly an order of magnitude from stage B to stage D, in line with the established Hyal-1/2/3-driven fragmentation model of CRC progression. EV-surface HA did not track this trajectory: EVs from stage B and stage D cells alike carried predominantly short, LMW-HA-dominant chains, and even EVs from HMW-HA-rich stage B cells were architecturally closer to stage D cells than to their own plasma membrane. This is not consistent with EV-surface HA being a passive membrane carryover during budding; it implies that EV biogenesis (or the endosomal sorting pathway that specifically gives rise to exosome-enriched EVs) actively selects for short HA chains, independent of the chain-length state prevailing at the cell surface at the moment of budding. The coarse-grained wrapping simulations offer a plausible physical basis for such a preference: short, sparsely grafted chains minimize the steric and bending-energy penalty of deforming a lipid bilayer around a polymer-decorated interface, a penalty that would apply whether the bilayer is deforming outward around a nascent vesicle or inward around an arriving one. We modelled the latter process explicitly, a target membrane wrapping an HA-decorated vesicle, because it is the process that the simulation geometry and the subsequent uptake experiments are built to test. Whether the same energetic logic also governs the reverse (budding) process during biogenesis is a reasonable physical inference, but was not directly simulated here and merits its own dedicated model.

### The uptake data initially appear to contradict the density prediction, and resolving that tension is instructive

Native HCT116 EVs carry the highest HA density of any group (∼17.9%) yet are also the most efficiently internalized, the opposite of what density alone would predict from **Fig. 3C**. The resolution lies in chain length: Mod-HCT116 EVs, despite lower density (13.1%), acquire a long HA population (∼4361 nm) entirely absent from any native sample, and the (L, φ_s_) phase map shows wrapping efficiency is substantially more sensitive to chain-length than to density in this regime. The uptake data are therefore consistent with the simulation once both parameters, not density in isolation, are considered jointly, and they suggest chain length is the dominant regulator of EV internalization efficiency in this system. A direct quantitative overlay of the measured (density, L_c_) coordinates for each EV population onto the **Fig. 3E** phase map would make this argument considerably stronger.

### HA density and CD44 expression are dissociated on EVs, which argues against a simple receptor-density explanation

Cellular CD44 levels increased with stage, in parallel with HA density, consistent with the established CD44-HAS autocrine loop. EV CD44 did not differ significantly by stage despite a large EV HA density difference, indicating that EV-surface HA loading is not simply tracking bulk CD44 abundance and is governed by a distinct, presumably biogenesis-intrinsic, mechanism. This dissociation strengthens the case that EV HA architecture is an independent regulatory axis rather than a downstream readout of the same program that elevates cell-surface HA.

### MVs and small EVs respond differently to HMW-HA reprogramming, pointing to distinct sorting mechanisms across biogenesis routes

HMW-HA supplementation substantially remodeled small-EV HA density, stiffness, and (partially) chain length, but left MV HA architecture essentially unchanged. Because MVs bud directly from the plasma membrane while exosome-enriched EVs are generated through the endosomal/multivesicular body pathway, this asymmetry suggests that active, selective HA sorting is a feature of the endosomal route specifically, while MV composition more directly and rigidly mirrors bulk plasma membrane state at the moment of budding. This distinction, if it holds, has practical consequences for biomarker design: small-EV HA architecture appears to be the more dynamic and therapeutically engineerable compartment, while MV HA architecture may be the more stable indicator of baseline cell-surface state.

### Limitations

All measurements were performed on established CRC cell lines rather than on primary patient tumors, ascites, or plasma-derived EVs, so the absolute density and chain-length values reported here should be treated as a proof-of-principle-stage signature. The coarse-grained model captures generic polymer-brush/membrane energetics but does not include receptor-mediated adhesion (e.g., CD44-HA bonds), bilayer compositional heterogeneity, or active cytoskeletal remodeling during real endocytosis, any of which could shift the balance between density and chain-length sensitivity observed here. Pharmacological (4-mU) and enzymatic (hyaluronidase) perturbations, while consistent and dose-graded, are not genetic loss-of-function approaches (e.g., HAS2/HAS3 or HYAL knockdown) and could in principle have off-target effects on other glycosaminoglycans. Finally, the isogenic and mechanistic-perturbation experiments were performed in a subset of lines (SW480/SW620) rather than across the full panel, and stage C is represented by a single line.

### Significance

These results reframe tumor HA not as a single overexpressed biomarker but as an architecturally dynamic system with at least two independently regulated compartments, a fragmenting cell surface and a consistently short-chain-enriched EV surface, that are mechanically contributory, functionally consequential for intercellular communication, and experimentally reversible. This positions HA surface architecture, and specifically EV-surface HA chain length and density, as a candidate for both a stage-resolving liquid-biopsy signature and a druggable node for limiting EV-mediated tumor communication in advanced CRC, motivating validation in patient-derived material as the next step.

## Materials and Methods

### Materials

L-15 (AL011A), DMEM (AL294A), RPMI-1640 (AL028), McCoy’s 5A (AL057H), EMEM (AL047), antibiotic solution (A001A), and PBS (ML116) were purchased from HiMedia, India. EV-free FBS (A2720801), CO₂-independent medium (18045088), CD63-FITC monoclonal antibody (MEM-259), CD9 monoclonal antibody (MA5-31980), goat anti-rabbit IgG FITC secondary antibody (64-6111), streptavidin Alexa Fluor™ 647 (S21374), and aldehyde/sulfate latex beads (4% w/v, 4 µm; A37304) were obtained from Thermo Fisher Scientific, USA. TEM grids (930253), biotinylated hyaluronic acid-binding protein (385911), hyaluronidase (H3506), DAPI (10236276001), (3-aminopropyl)triethoxysilane (440140), 4-methylumbelliferone (M1381), and polystyrene microparticles (79633) were purchased from Sigma-Aldrich, USA. APC anti-human CD81 (TAPA-1) antibody was obtained from BioLegend, USA. MAL-dPEG®₂₄-NHS ester was purchased from Vector Laboratories, USA. Doxorubicin (D4193) was purchased from TCI, Japan. Hyaluronic acid fluorescein (HA-804) was obtained from Creative PEGWorks, USA. Hyaluronic acid sodium salt (75810) and uranyl acetate dihydrate (81405) were purchased from SRL, India. All reagents were used as received without further purification.

### Cell culture

CRC cell lines SW480, HT-29, SW620, Colo-205, and HCT116 were obtained from the National Centre for Cell Science (NCCS, India), while the non-malignant human colon fibroblast cell line CCD-18Co was obtained from the American Type Culture Collection (ATCC, USA). SW480 and HT29 represent Dukes’ stage B, SW620 stage C, and HCT116 and COLO205 stage D, as previously reported^33,34^. Cells were cultured according to the suppliers’ protocols in their respective media (L-15, DMEM, RPMI-1640, McCoy’s 5A, and EMEM) supplemented with 10% EV-depleted FBS and 1% antibiotics, and maintained at 37 °C with 5% CO₂ until reaching 80–90% confluence before conditioned media were collected for EV isolation.

### 4mU and HMW-HA treatment

HCT116 cells were cultured in ten T75 flasks using the complete medium described above. At the desired confluency, the medium was replaced with FBS-free medium containing either 1 mM 4-methylumbelliferone (4-MU) or 250 µg/mL HMW-HA (1500 kDa) and incubated for 24 h. The treatment medium was then replaced with fresh FBS-free medium, followed by an additional 24–48 h incubation before conditioned media were collected for EV isolation^35–38^.

### EV and MV isolation

EVs were isolated by differential ultracentrifugation according to previously reported protocols and the Minimal Information for Studies of Extracellular Vesicles (MISEV) guidelines^23^. Conditioned media were sequentially centrifuged at 500 × g for 10 min, 3,000 × g for 15 min, and 16,500 × g for 30 min (4 °C) to remove cells, debris, and isolate microvesicles (MVs). The MV pellet was washed with PBS, recentrifuged at 16,500 × g for 30 min, resuspended in PBS, and stored at −80 °C. The EV-containing supernatant was filtered through a 0.22 μm membrane and ultracentrifuged at 120,000 × g for 70 min at 4 °C using an SW-41Ti rotor (Beckman Coulter, USA). The EV pellet was resuspended in PBS and stored at −20 °C for short-term or −80 °C for long-term use.

### Nanoparticle tracking analysis

EV concentration and size distribution were analysed by nanoparticle tracking analysis (NTA) using a NanoSight NS500 system (Malvern Instruments) equipped with a 532 nm laser at 20 ± 3 °C. EV samples were diluted to achieve an optimal concentration of 20–100 particles per frame, and each sample was measured five times. The recorded videos were used for particle counting, and size distribution data were analysed using ZetaView software.

### Transmission electron microscopy

EV samples were deposited onto glow-discharged, carbon-coated copper grids for 30 s, blotted, and negatively stained with 2% (w/v) filtered uranyl acetate for 30 s. After air-drying, samples were imaged using a JEM-F200 (CF-HR) transmission electron microscope (JEOL Ltd., Japan) operated at 200 kV with a probe current of 2.5 nA and a spatial resolution of 0.7 nm.

### Flow cytometry

EVs and MVs were characterised for CD9, CD63, CD81, and HA using a bead-based flow cytometry assay. Briefly, EVs were adsorbed onto 4-µm aldehyde/sulfate latex beads, incubated overnight at 4 °C, and stained with anti-CD9, FITC-conjugated anti-CD63, APC-conjugated anti-CD81, or biotinylated hyaluronic acid-binding protein (HABP; 5 µg/mL). For HABP staining, streptavidin–Alexa Fluor 647 (1:200) was used, while CD9 was detected using goat anti-rabbit IgG FITC secondary antibody (1:200). Following washing, bead–EV complexes were analysed on a Beckman Coulter CytoFLEX S flow cytometer, and data were processed using Floreada.io software.

For HA removal, EVs were treated with hyaluronidase (10 U/mL, 1 h) before bead capture and HA staining. Cell-surface HA was analysed by staining trypsinised cells with biotinylated HABP (5 µg/mL), followed by streptavidin–Alexa Fluor 647. For HA digestion, confluent cells were treated with hyaluronidase (10 U/mL, 1 h at 37 °C) before staining and quantification, as previously described^39^.

### Confocal Microscopy

Cells were cultured on 35-mm glass-bottom dishes, washed with PBS, and stained with 1× CellMask Green for 10 min at 37 °C before fixation with 4% paraformaldehyde (PFA) for 20 min. For HA removal, cells were treated with hyaluronidase (10 U/mL) for 10 min before fixation. Fixed cells were incubated with biotinylated hyaluronic acid-binding protein (HABP) overnight (16 h), followed by Alexa Fluor 647-conjugated streptavidin (1:200) for 1 h in the dark. All samples were counterstained with DAPI (1 µg/mL), and fluorescence images were acquired using a Nikon AXR NSPARC laser-scanning confocal microscope (Ti2E, 100× Apochromat oil immersion objective, NA 1.45).

### Atomic Force Microscopy

Cells were cultured on sterile glass coverslips in their respective complete culture media supplemented with 10% fetal bovine serum (FBS) at 37 °C in a humidified atmosphere containing 5% CO₂ for 24–48 h. Upon reaching the desired confluency, the cells were fixed with 4% paraformaldehyde for 10 min, thoroughly washed with PBS, and stored in PBS until AFM analysis. For EV imaging, freshly cleaved muscovite mica substrates (Ted Pella, USA) were mounted onto clean glass slides using adhesive. Freshly isolated EVs were deposited onto the mica surface (30 µL per sample) and allowed to adsorb for 10 min at room temperature (25 °C). The substrates were subsequently rinsed three times with 200 µL PBS to remove loosely adhered vesicles and maintained in PBS while imaging. Topographical imaging of cells and EVs was performed in PBS using a MFP-3D Origin AFM, Asylum Research, Oxford Instruments. Cells were imaged in contact mode using an MLCT-C probe (Bruker, Santa Barbara, CA, USA) spring constant ∼0.01 N m⁻¹. EVs were imaged in tapping mode using an SNL-10 D probe, spring constant ∼0.1 N m⁻¹ (Bruker, Santa Barbara, CA, USA). All measurements were carried out at room temperature. The acquired AFM topographical images were processed using Igor Pro software. Dimensional analysis of individual EVs was subsequently performed to determine their diameter, height, surface area, and surface roughness from the AFM images.

To quantify the surface density of HA on both cell and EV membranes, HABP functionalized AFM probes were prepared according to previously established protocols^20^. Briefly, SNL-10 D cantilevers were sequentially cleaned with ultrapure water and ethanol, followed by drying under a nitrogen stream. The probes were amine-functionalized by incubating them in a 5% (v/v) solution of (3-aminopropyl) triethoxysilane (3-APTES) prepared in a 5:95 ethanol–water mixture. An aldehyde-PEG-NHS linker was then conjugated to the cantilever tips by incubating the probes for 2 h in a reaction mixture containing 500 µL chloroform, 30 µL triethylamine, and 1 mg Maleimide-PEG-NHS. Following linker attachment, the probes were washed three times with chloroform, dried under nitrogen, and incubated with 20 nM HABP for 30 min at room temperature in a sealed Petri dish. The functionalized probes were rinsed with ultrapure water immediately before use for AFM-based single-molecule force spectroscopy measurements.

Force distance curves were collected at a typical velocity of 0.5 µm·s⁻¹; for loading rate related studies, the retraction velocities were varied between 1.5 and 10 µm·s⁻¹. The maximum applied force was limited to 650 pN, and the surface dwell time at peak load was kept as short as possible. The cantilever spring constant and deflection sensitivity were calibrated using the thermal tune method, always prior to starting any force spectroscopy experiment. All measurements were repeated at least twice using independently prepared samples with functionalized AFM probes (**Table S2**). Force data were analyzed using Igor Pro data processing software (Asylum Research). Force maps were acquired over the nuclear regions of fixed cells at a scan rate of 1 Hz and a resolution of 32 × 32 pixels across a 250 nm × 250 nm area. Individual EVs were initially imaged within scan areas of 200 × 200 nm², then single vesicles were further zoomed into an 80 × 80 nm² scan area. Force maps were then recorded over the central region of the zoomed EVs at a resolution of 32 × 32 pixels across a 50 nm × 50 nm area. Based on the specific adhesion forces corresponding to HA–HABP interactions^20^, a predefined force threshold was applied to convert the gradient force maps into binary, color-coded maps, where specific interactions were denoted in green and nonspecific or negative interactions in black. Automated pixel counting was then performed to quantify positive and negative interaction events, enabling estimation of HA surface density on cells and EV surfaces. The interaction between HABP and HA exhibited a linear fit to the, Bell–Evans model^40^. Which describes a linear dependence of the unbinding force on the logarithm of the loading rate, as expressed in Equation 2.

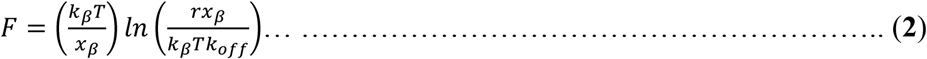

Where *F* represents the most probable unbinding force, *x*_β_ denotes the distance from the bound state to the transition state, *r* is the loading rate calculated as the product of the cantilever retraction velocity and its effective spring constant, and *k*ₒ_ff_ corresponds to the kinetic off-rate constant.

The contour lengths of individual HA chains were determined by fitting the force–separation curves using the worm-like chain (WLC) model with FODIS software^41^ Within this framework, each unfolding event is described by Equation 3.

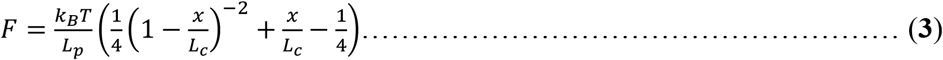

Here, *F* denotes the applied force and *x* represents the extension; *k*_B_ is the Boltzmann constant, *T* is the absolute temperature, *L_p_ (*varied between 4.1 and 4.4 nm) corresponds to the persistence length, and *L*_c_ is the contour length of the domain^21^.

For AFM nanoindentation and viscosity measurements, commercial tipless cantilevers (HQ:NSC36 C, Micromasch, Bulgaria) were used. A 5µm glass bead was attached to the tipless cantilever following a previously published protocol^42^. The spherical probe was operated in approach-dwell-retract cycles on single cells to simultaneously evaluate their elastic and viscous responses. During these cycles, force curves, including relaxation profiles, were recorded. Single force–indentation curves and force-volume maps were acquired using a maximum applied force of 300 pN for EVs and 1–2 nN for cells. For each condition, measurements were performed on fresh samples prepared independently three times. Young’s modulus values were determined from the force curves using the Hertz model (Equation 4)

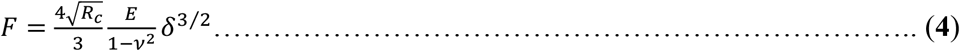

Here, δ denotes the indentation depth, ν represents the Poisson’s ratio, R_c_ corresponds to the cantilever tip radius, and E is the elastic modulus. The Poisson’s ratio was taken as 0.3^21^.

The relaxation time of cells was calculated from the relaxation curves by applying 2^nd^ order Maxwell spring-dashpot model (Equation 5).

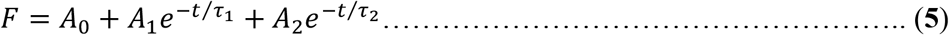

A₀ represents the instantaneous response, A₁ and A₂ denote the force amplitudes, τ_1,_ and τ_2_ are the fast and slow cellular relaxation time respectively. The relaxation curves were fitted using the Maxwell model with a custom program implemented in Origin Pro software. Following the determination of the cellular Young’s modulus using the Hertz model and the relaxation time using the Maxwell model, the cellular viscosity was subsequently calculated (Equation 6).

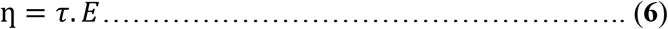

Here, η denotes the cellular viscosity, τ represents the cellular relaxation time, and E corresponds to the cellular Young’s modulus^43^.

### Theoretical model and methods

We study the wrapping of a polymer-grafted vesicle by a fluid membrane using coarse-grained molecular dynamics simulations performed with LAMMPS^44^. The system has three components. The membrane is a one-particle-thick, solvent-free fluid sheet described by the anisotropic pair potential of Yuan et al.^45^, in which each particle represents a cluster of lipid molecules and is characterized by its center-of-mass position ri and a unit orientation vector ni. In the model, the interaction between two membrane particles comprises a soft-core repulsive term and an attractive term whose strength depends on the relative orientations of the particles with respect to their separation vector. The explicit form of the anisotropic interaction potential is given in the original reference. The parameters governing the membrane properties, including bending rigidity, in-plane fluidity, and spontaneous curvature, along with their respective values, are summarized in **Table S3**. The vesicle is modeled as a spherical shell of beads with radius R=10σ, treated as a single rigid body with a fixed shape; all vesicle beads undergo a common translation and rotation. Semiflexible bead–spring polymer chains are grafted onto the vesicle surface and modeled using harmonic bond and cosine angle potentials to control their connectivity and stiffness. At the grafting points, enhanced bond and angle stiffness are employed to constrain the chains to emerge approximately normal to the vesicle surface. The chain length L and the grafting density φ_s (the fraction of vesicle surface sites carrying a chain) are the two control parameters of this study.

All non-membrane pairs interact through a Lennard–Jones (LJ) potential. Polymer–polymer, membrane–vesicle, and vesicle–polymer interactions are modeled as purely repulsive Weeks–Chandler–Andersen (WCA) interactions with a cutoff r_c = 2^(1/6)^σ. In contrast, the polymer–membrane interaction responsible for membrane wrapping is modeled using the full LJ potential, with a well depth ɛ_pm_=2.2 k_B_T and a cutoff r_c_=2.5 σ. All quantities are expressed in reduced Lennard–Jones units, where (m), , (k_B_T), and τ) denote the monomer mass, characteristic particle diameter, thermal energy, and time unit, respectively. The equations of motion are integrated using Langevin dynamics in LAMMPS. The membrane and polymer monomers are coupled to a Langevin heat bath, whereas the vesicle is integrated as a single rigid body and is not directly thermostatted. Each system undergoes an initial relaxation, followed by gradual heating and equilibration to the target temperature. Production simulations are subsequently performed at k_B_T=1.0 with a timestep of **Δt=3×10^−4^τ** and are continued until the wrapping fraction reaches a steady state.

### Visualizing HMW-HA on stage D cell surface

HCT116 cells (1 × 10⁵) were seeded on 35-mm glass-bottom dishes and cultured in complete McCoy’s 5A medium for 24–48 h. Cells were then incubated with 5 µg/mL biotinylated HABP in FBS-free medium for 16 h, followed by streptavidin–Alexa Fluor 647 (1:200) for 1 h. After washing, cells were incubated with 250 µg/mL FITC-conjugated HMW-HA (1500 kDa) in FBS-free medium for 24 or 48 h. Cells were then washed, fixed with 4% paraformaldehyde for 15 min, counterstained with DAPI (1 µg/mL), and imaged using a Nikon AXR NSPARC laser-scanning confocal microscope (Ti2-E) with 20× and 100× objectives. For quantitative analysis, cells were processed using the same protocol, trypsinised after FITC-HMW-HA incubation, washed, and analysed by FACS.

### Wound Healing Assay

Cell migration was evaluated using a scratch (wound-healing) assay. HT29 and HCT116 cells (1 × 10⁶ cells/well) were seeded in 6-well plates and grown to ∼90% confluence. A uniform scratch was created using a sterile 200 µL pipette tip, and detached cells were removed by washing with PBS. HCT116 cells were then cultured in serum-free McCoy’s 5A medium with or without 1 mM 4-MU, 10 U/mL hyaluronidase, or 250 µg/mL HMW-HA, while HT29 cells were maintained in serum-free DMEM. Wound closure was monitored for 96 h, with images acquired at 0–96 h using a Nikon Eclipse TS2 microscope and analysed with ImageJ (NIH, USA). Experiments were performed in duplicate and repeated at least three times.

### EV Uptake

Cellular uptake of EVs was evaluated by confocal microscopy and flow cytometry. Cells were cultured in glass-bottom or plastic dishes and, at ∼80% confluence, were maintained in FBS-free medium with or without 250 µg/mL HMW-HA for 24 h. Purified EVs (100 µg) were loaded with doxorubicin (DOX; 50 µg) by electroporation (350 V, 150 µF) in 200 µL electroporation buffer using a Gene Pulser II Electroporator (Bio-Rad, USA). Free DOX was removed using 100 kDa Amicon filters (4500 × g, 15 min), and encapsulated DOX was quantified using a plate reader (λ_ex = 490 nm, λ_em = 590 nm). DOX-labelled EVs were incubated with recipient cells for 6 h at 37 °C, followed by PBS washing, fixation with 4% paraformaldehyde, and DAPI (1 µg/mL) staining. EV uptake was visualised using a Nikon AXR NSPARC laser-scanning confocal microscope (Ti2-E, 100× objective) and quantified by flow cytometry.

## Data availability

The data reported in this study are available within the article and its Supplementary Materials file. Raw data are available from the corresponding authors upon reasonable request.

## Acknowledgements

The authors thank Shiv Nadar Institution of Eminence for research facilities, including MFP-3D AFM. The authors acknowledge the DST-FIST grant (SR/FST/LS-1/2017/59(c)) for the confocal microscopy facility at Shiv Nadar Institution of Eminence. We thank Prof. Sanjeev Galande’s lab for providing the SW620 cell line.

## Author contributions

D.P.: Investigation, methodology, data analysis, writing-review, and editing. T.A. and N.N.: Investigation. N.M. and A.K.D.: Computational study, methodology, and data analysis; writing methodology and results. S.S: Investigation. S.P.: Funding acquisition, writing-review and editing. T.R.: Conceptualization, supervision, writing-review and editing, and funding acquisition.

## Additional information

Data which supports this study provided in the supporting information

**Correspondence** and requests for materials should be addressed to

## Supporting Information

**Figure S1.**
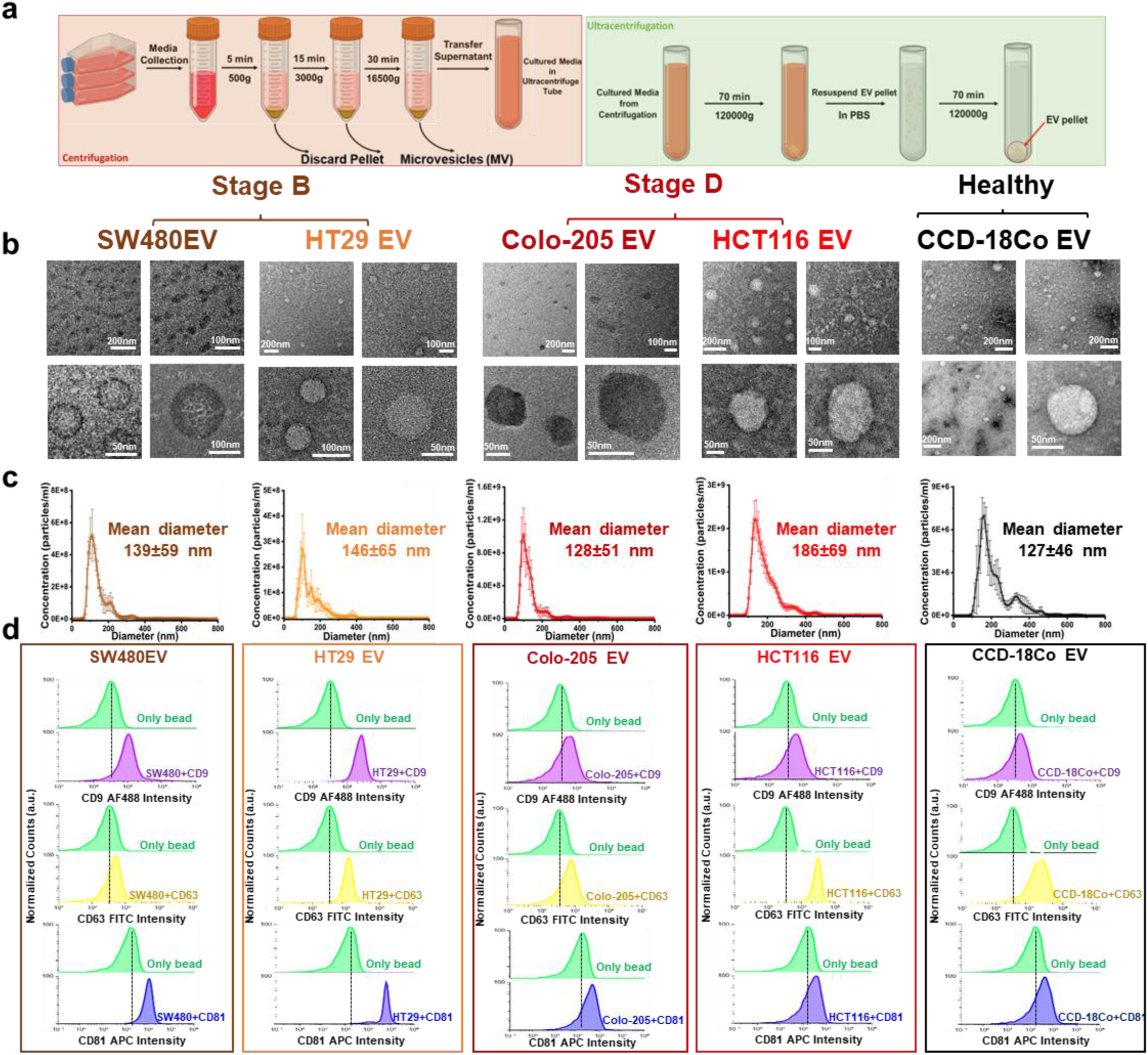
EV isolation and characterization. **(a)** Schematic illustration of the differential ultracentrifugation workflow employed to isolate EVs from cell culture conditioned media. **(b)** Representative transmission electron microscopy (TEM) images showing the near spherical morphology of EVs derived from CRC & healthy cells. **(c)** Nanoparticle tracking analysis (NTA) based size distribution profiles of EVs. **(d)** Flow cytometry analysis showing enrichment of tetraspanin markers CD9, CD63, and CD81 in CRC-derived & healthy EVs.

**Figure S2:**
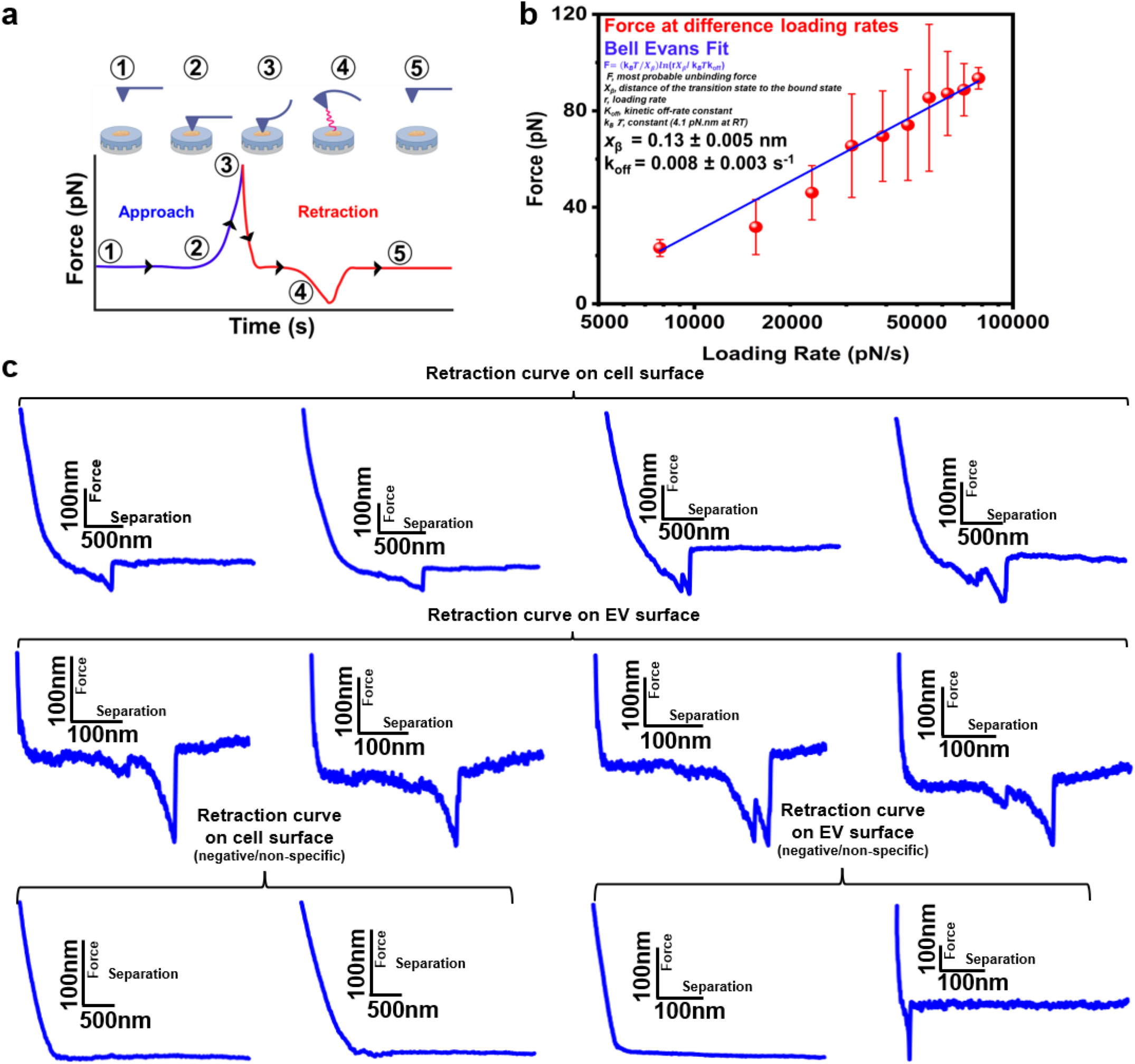
AFM Unbinding experiments, Bell–Evans model. **(a) Schematic illustration of the SMFS experimental setup, showing an AFM cantilever functionalized with HABP probing HA molecules on the surface of CRC cells & EVs.** (b) Dynamic force spectroscopy analysis of HABP–HA bond dissociation, with rupture force plotted as a function of loading rate and fitted to the Bell–Evans model (blue solid lines); shaded regions represent the 95% confidence intervals, confirming specific interactions. (c) Representative force-distance (retract) curves obtained, including specific adhesion events, non-specific interactions, and no-adhesion events, were recorded on both CRC cells and EVs derived from different disease stages.

**Figure S3:**
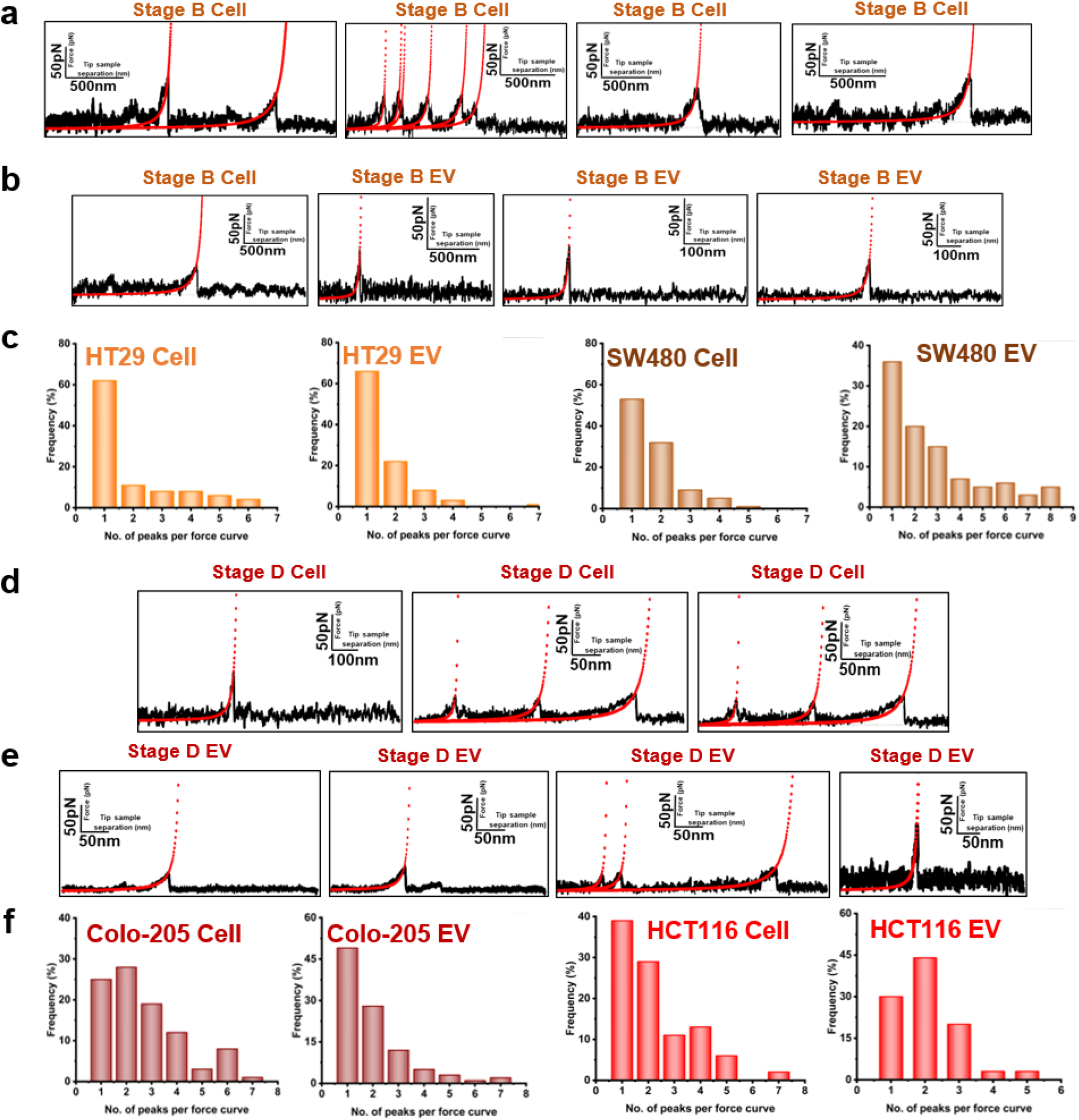
Analysis of SMFS curves of specific HA–HABP interactions, worm-like chain (WLC) model (a, b) Representative retract-force–distance curves (black traces) acquired on the surfaces of stage B CRC cells and their EVs, respectively, showing single and multiple HA chain stretching and unbinding events. The corresponding worm-like chain (WLC) model fits are overlaid in red, (c) Histogram of the distribution of rupture events’ number per individual force–distance curve obtained from stage B CRC cells and EVs, reflecting the relative contributions of single- and multi-interactions. (d, e) Representative retract-force–distance curves (black traces) recorded on stage D CRC cells and their EVs, respectively, demonstrating mono- and multi-chain HA stretching and unbinding signatures; WLC model fits are shown in red. (f) Histogram of the number of rupture events per force–distance curve obtained from stage D CRC cells and EVs, highlighting the increased prevalence of multiple unbinding events associated with advanced disease stage.

### System construction and simulation details for the diffusion calculation

To understand how the grafted polymers affect the mobility of the vesicle, we performed a separate set of simulations in which a polymer-grafted vesicle diffuses freely in a simulation box, in the absence of the membrane. We use the same rigid spherical vesicle (*R* = 10*σ*) as in the main text, onto which the polymer chains are grafted; the chains are described by the same harmonic bonds and cosine-angle stiffness used there. For the diffusion study we varied the polymer length over *L* = 10-40*σ* and the surface density (the fraction of vesicle surface sites carrying a chain) over *ϕ*_s_ = 0.05-0.50. Here, polymer–polymer and vesicle–polymer pairs interact through the Weeks–Chandler–Andersen (WCA) potential, that is, a Lennard-Jones potential with *ε* = 1 *k*_B_*T* and *σ* = 1 truncated at its minimum, *r*_c_ = 2^1/6^*σ*. The vesicle is treated as a rigid body and the grafted beads follow Newtonian dynamics, with both coupled to a Langevin thermostat at *k*_B_*T* = 1. Each system is first equilibrated and then run in production for a sufficiently long time to span both the short-time ballistic and the long-time diffusive regimes.

#### Mean-squared displacement and diffusion coefficient

From the production trajectory we compute the mean-squared displacement (MSD) of the grafted chains,

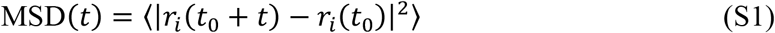

averaged over all grafted beads *i* and over time origins *t*_0_. At short lag times the motion is ballistic (MSD ∼ *t*^2^) and crosses over at long lag times to the diffusive regime (MSD ∼ *t*), from which the long-time diffusion coefficient is obtained as

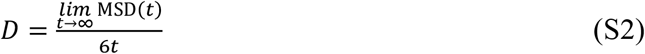

The resulting MSD curves and the extracted diffusion coefficients are shown in Figure S4.

**Figure S4:**
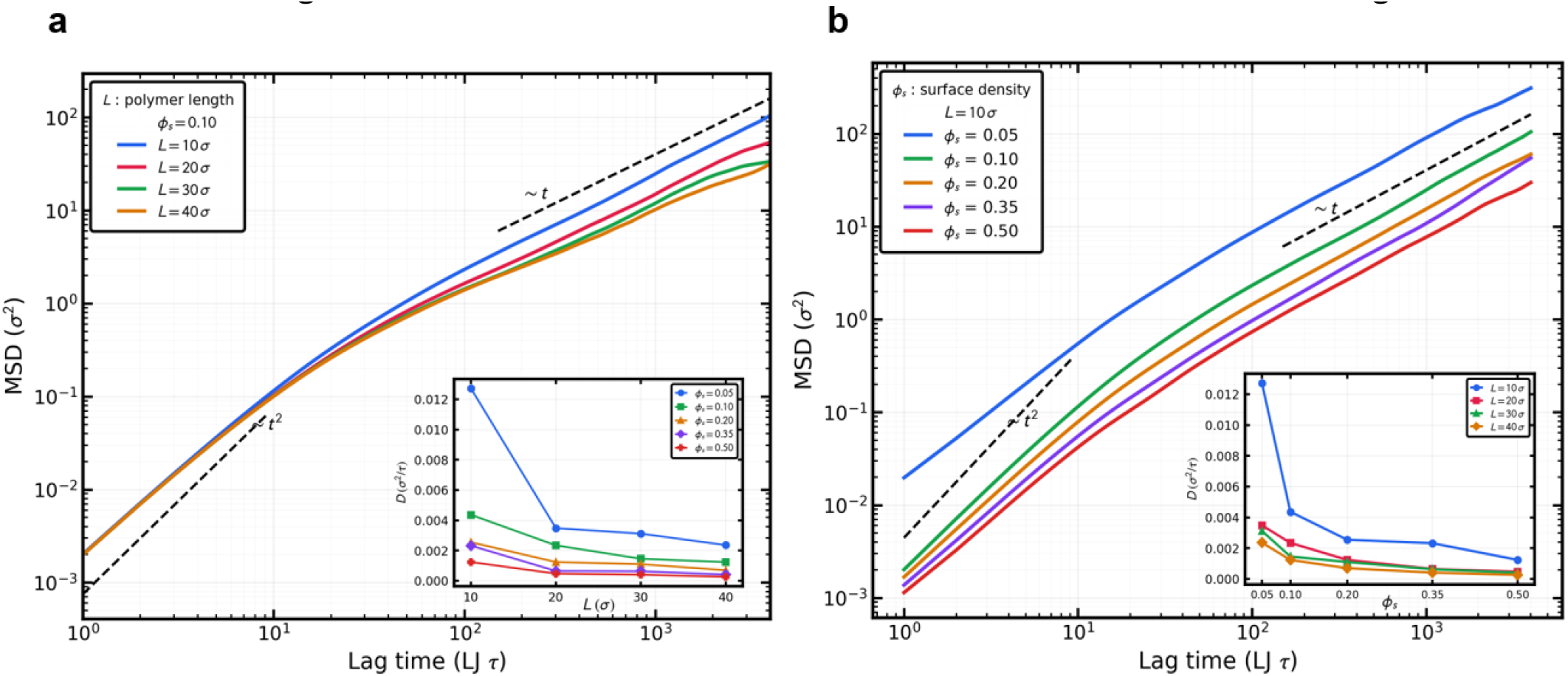
(a) MSD versus lag time for surface densities ϕ_s_ = 0.05-0.50 at fixed polymer length L = 10σ. Inset: diffusion coefficient D versus ϕ_s_ for several polymer lengths L. (b) MSD versus lag time for polymer lengths L = 10-40σ at fixed surface density ϕ_s_ = 0.10. Inset: diffusion coefficient D versus L for several surface densities ϕ_s_.

**Figure S5a:**
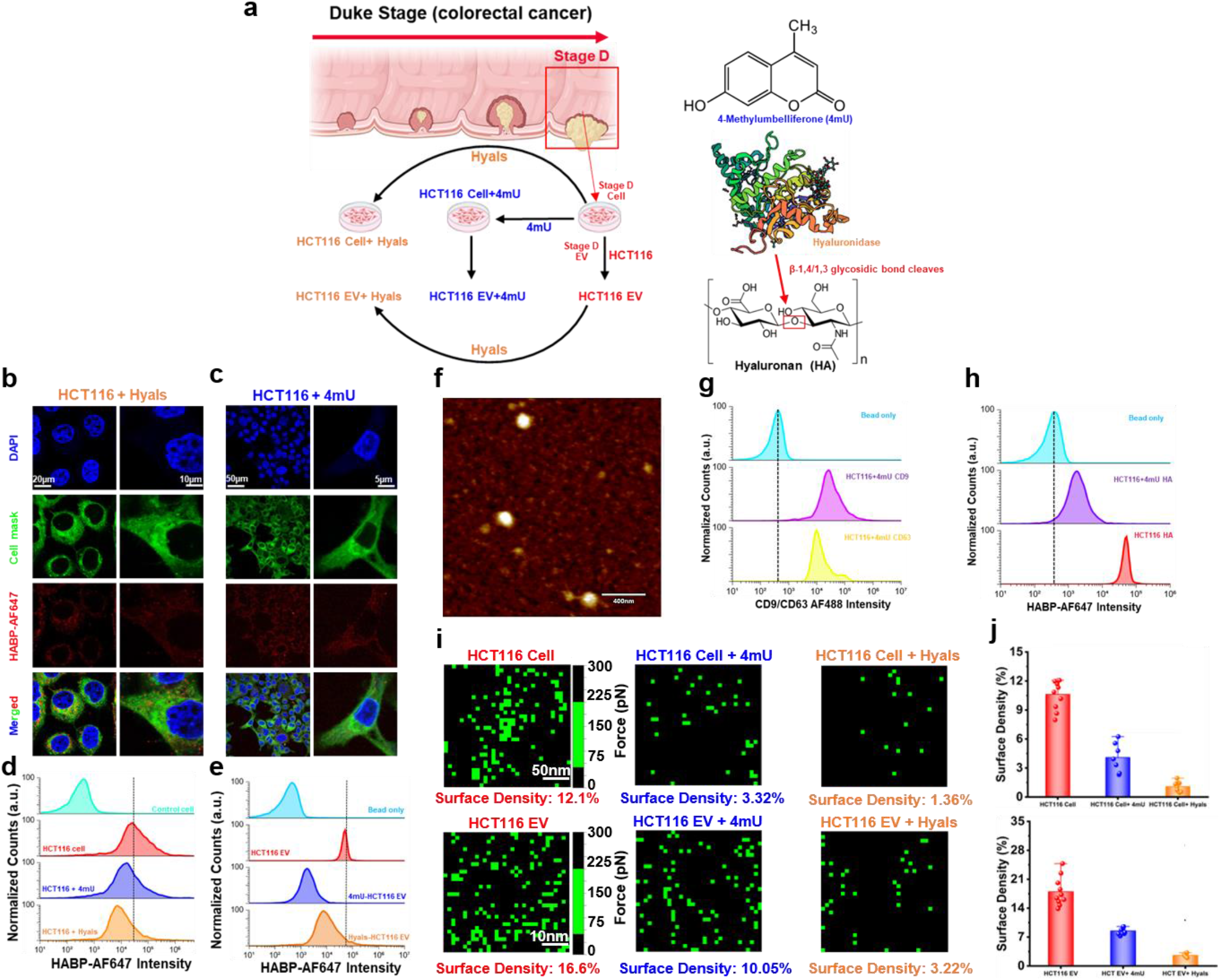
Modulating HA expression on HCT116 cells and EVs using 4 MU. **(a) Schematic depicting the experimental workflow in which HCT116 cells with 4-methylumbelliferone (4-MU) to inhibit HA biosynthesis or with hyaluronidase (Hyals) to enzymatically degrade surface HA and their EVs were prepared.** (b, c) Confocal microscopy images of 4-MU-treated cells, and hyaluronidase-treated cells stained with DAPI (blue) for nuclei, CellMask (green) for the plasma membrane, and HABP-AF647 (red) for surface HA. Merged images demonstrate a marked reduction in HA-associated fluorescence following both treatments. (d, e) Flow cytometry histograms showing quantitative analysis of HABP-AF647 fluorescence on intact HCT116 cells and EVs following HA modulation. Both 4-MU and hyaluronidase treatments significantly shift fluorescence intensity toward lower values relative to untreated controls, confirming efficient depletion of cell- and EV-associated HA. (f) Representative AFM topographical image showing isolated 4mU treated HCT116-derived EVs used for subsequent single-vesicle analyses. (g) Flow cytometry analysis of 4mU treated HCT116-derived EVs demonstrating preserved EV identity after 4-MU treatment through CD9/CD63-AF488 staining. (h) while HABP-AF647 fluorescence confirms a substantial reduction in EV-associated HA compared with untreated EVs. (i) SMFS-derived force maps illustrate the spatial distribution of HA-binding events on individual HCT116 cells (top panel) and EVs (bottom panel). Green pixels represent specific HA–HABP interaction events detected during force mapping, whereas black regions indicate the absence of detectable interactions. Quantitative analysis revealed a progressive reduction in HA surface density after treatment. (j) Bar graphs (right) summarize the quantitative HA surface density measurements obtained from SMFS for HCT116 cells (upper panel) and their corresponding EVs (lower panel), confirming that inhibition of HA biosynthesis and enzymatic degradation both significantly reduce HA abundance, with hyaluronidase producing the most pronounced depletion. Error bars represent the mean ± SD from independent biological replicates.

**Figure S5b:**
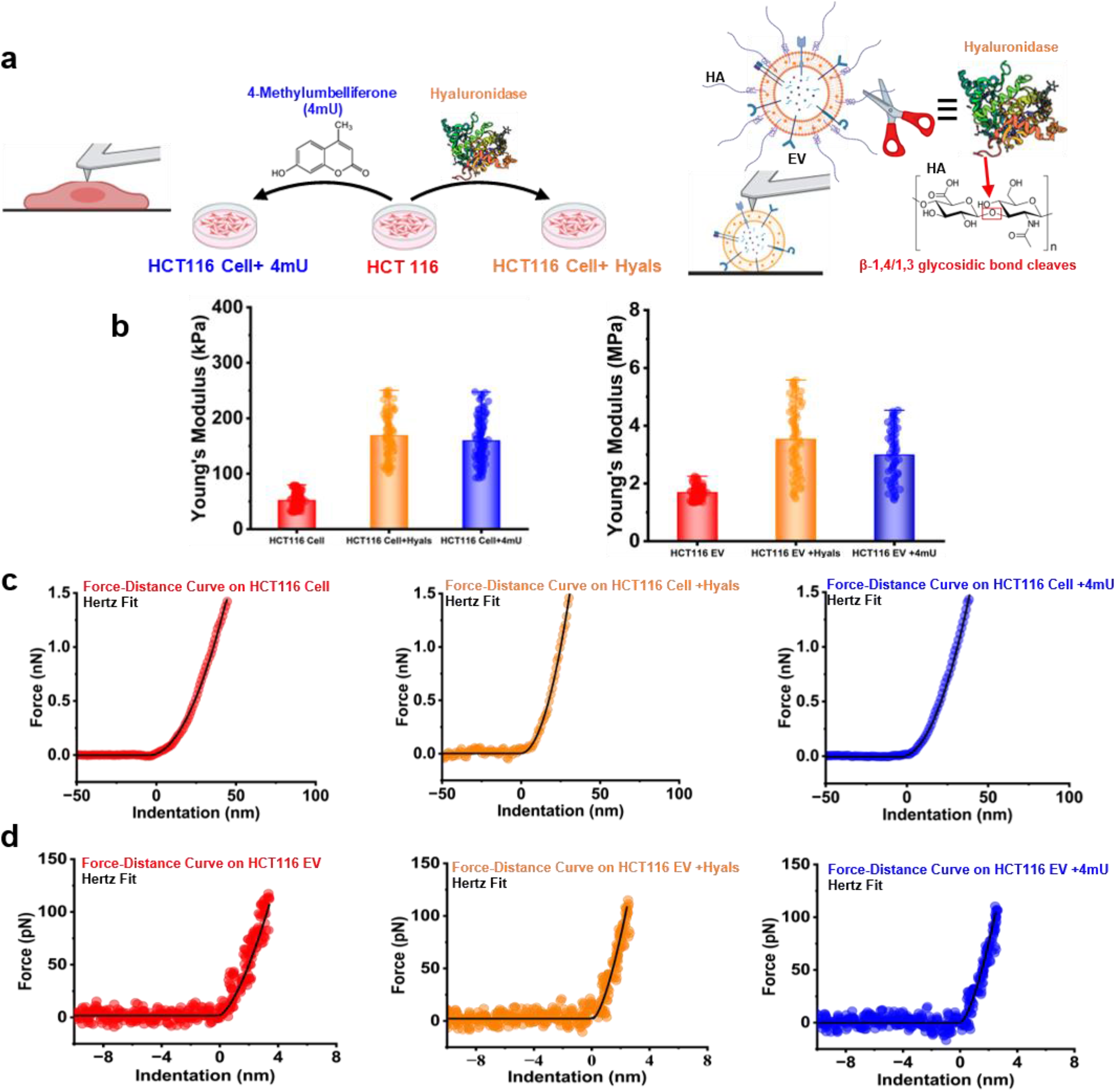
Effect of HA inhibition and degradation on the mechanical properties of stage D CRC cells and their EVs. (**a**) Schematic illustration of the enzymatic degradation of cell-surface HA by hyaluronidase treatment and 4-methylumbelliferone (4-mU) treatment on HCT116 (stage D) cells. (**b**) Cell and EV Young’s modulus before and after treatment. (**c, d**) Representative AFM force–indentation curves acquired from control (red), hyaluronidase-treated (orange), and 4-MU-treated (blue) stage D cells and their EVs. Solid black lines indicate Hertz model fits used to extract Young’s modulus values

**Figure S6:**
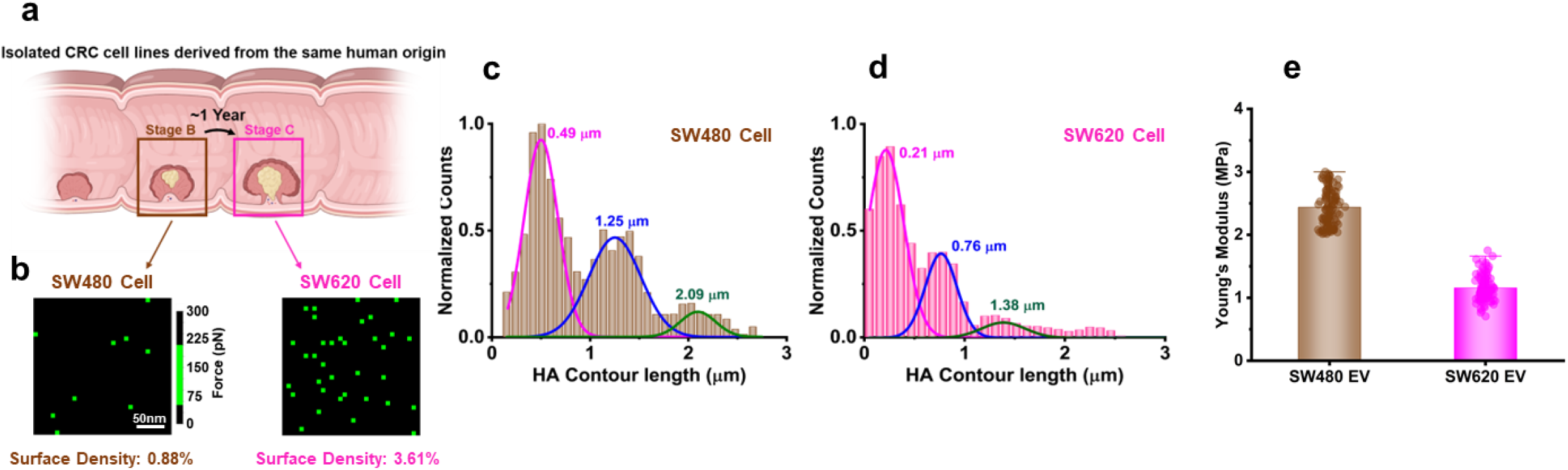
HA surface density and chain length distribution in CRC cells with same human source. (**a**) Schematic illustration of the experimental workflow for quantitative AFM-based analysis of HA surface density, along with representative force-volume adhesion maps acquired from CRC cell lines SW480 and SW620. (**b**) Quantitative comparison of HA surface density on SW480 and SW620 cells. (**c, d**) Distribution profiles of HA contour lengths (Lc) measured from stage B (SW480) and stage C (SW620) CRC cells, respectively. (**e**) Box plot of Young’s Modulus of the EVs.

**Figure S7:**
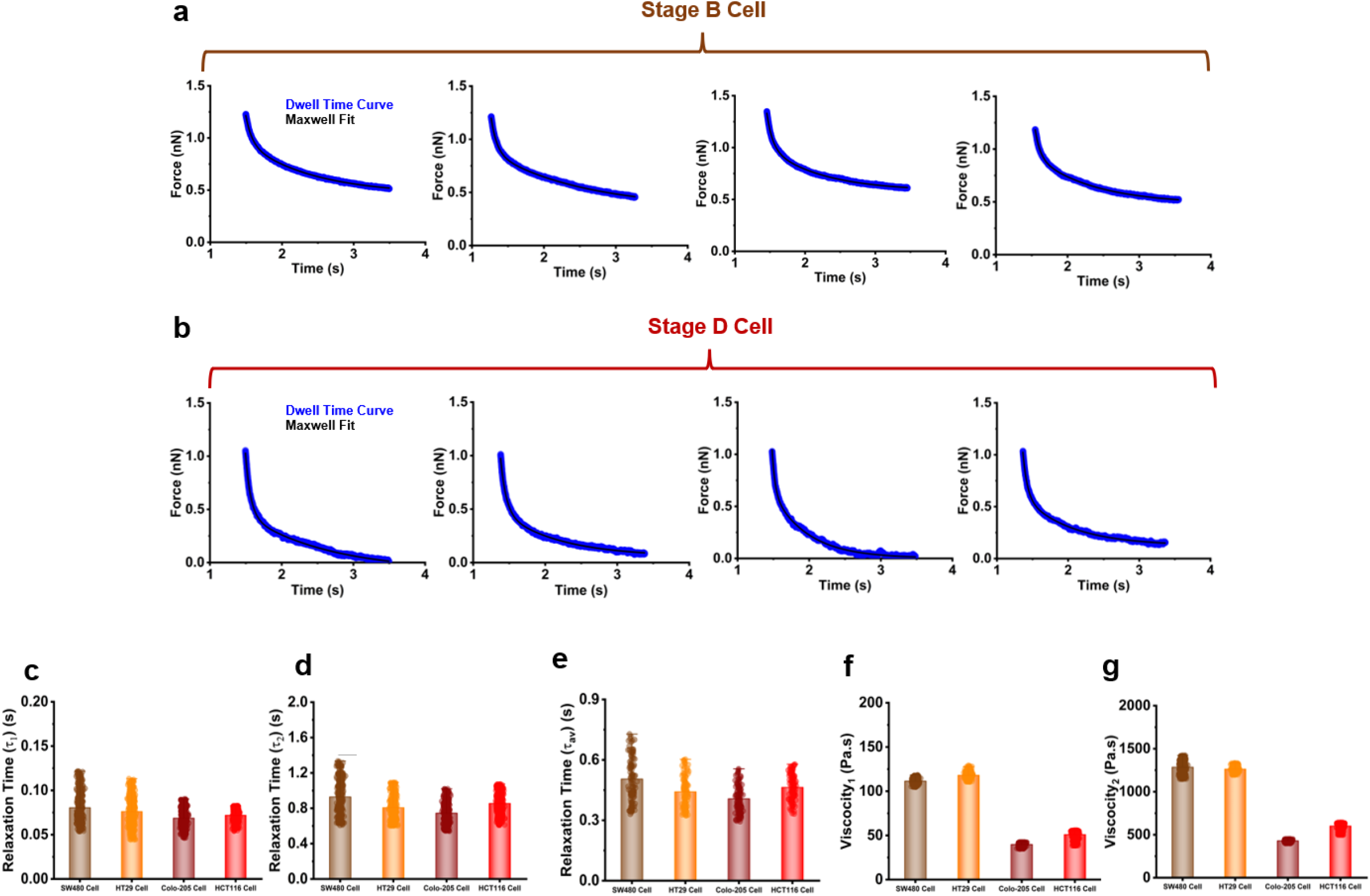
Stress-relaxation behavior and viscoelastic parameters of CRC cells across different disease stages. (a, b) Representative AFM-based stress-relaxation curves recorded on living stage B (a) and stage D (b) CRC cells. The experimental relaxation responses were fitted using a second-order (two-element) Maxwell viscoelastic model, shown as solid black lines, to extract characteristic cellular relaxation times. (c-e) Box plots summarizing the relaxation times τ₁, τ₂, and τ_av_ obtained from Maxwell model fitting of stress-relaxation curves measured on stage B and stage D CRC cells, respectively. (f, g) Box plots showing the apparent viscosities corresponding to the relaxation times τ₁ and τ₂ for CRC cells derived from different disease stages.

**Figure S8a:**
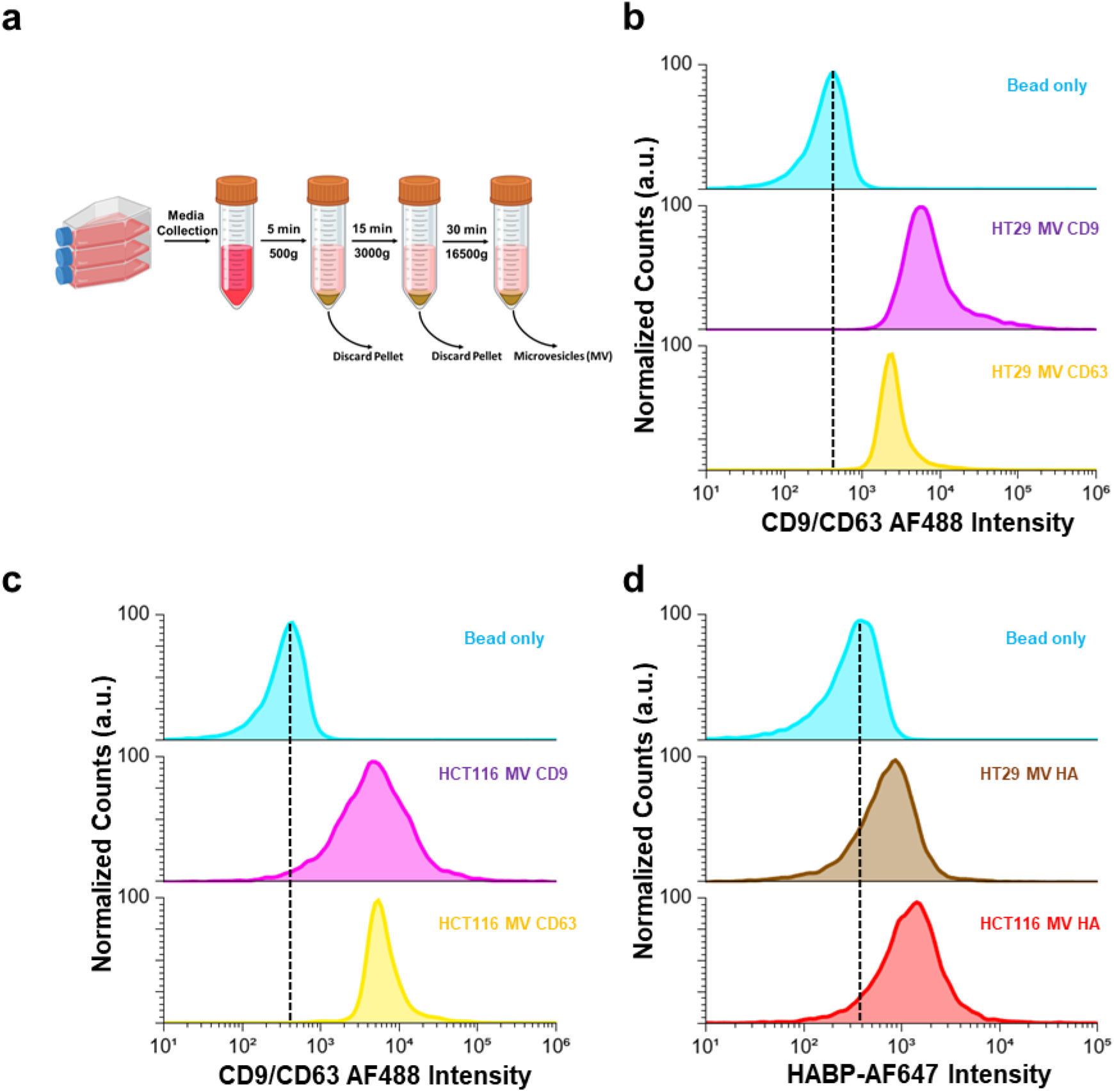
Isolation and characterization of MVs derived from stage B and stage D colorectal cancer (CRC) cells. **(a)** Schematic representation of the differential centrifugation workflow used to isolate MVs from cell culture conditioned media, **(b, c)** Flow cytometry analysis confirming tetraspanin markers CD9 and CD63 in MVs derived from stage B (HT-29) **(b)** and stage D (HCT116) (c) CRC cells. (d) Flow cytometry analysis showing the relative surface abundance of HA on CRC-MVs.

**Figure S8b:**
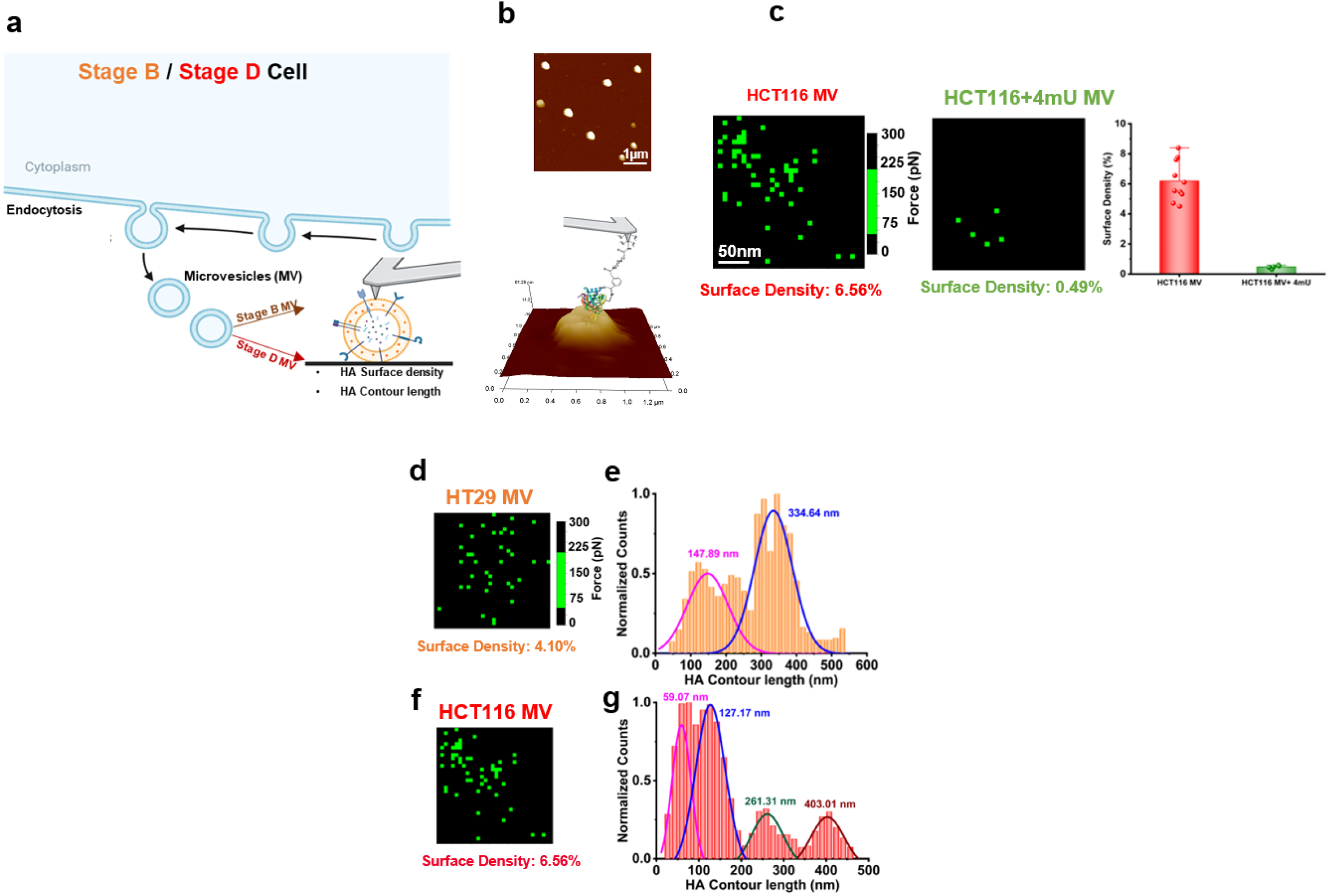
HA surface density and contour length on MVs, 4-MU treatment. (**a**) Schematic overview of the experimental workflow illustrating inhibition of HA synthesis in stage D CRC cells using 4-methylumbelliferone (4-mU), followed by isolation of MVs (**b**) Representative AFM image of MV, (**c**) unbinding force map showing the HA surface density measured on HCT 116 EVs and 4-MU treated MVs, corresponding quantitative box plot. (**d-g**) stage B and D CRC MVs’ HA surface density and contour lengths.

**Figure S9:**
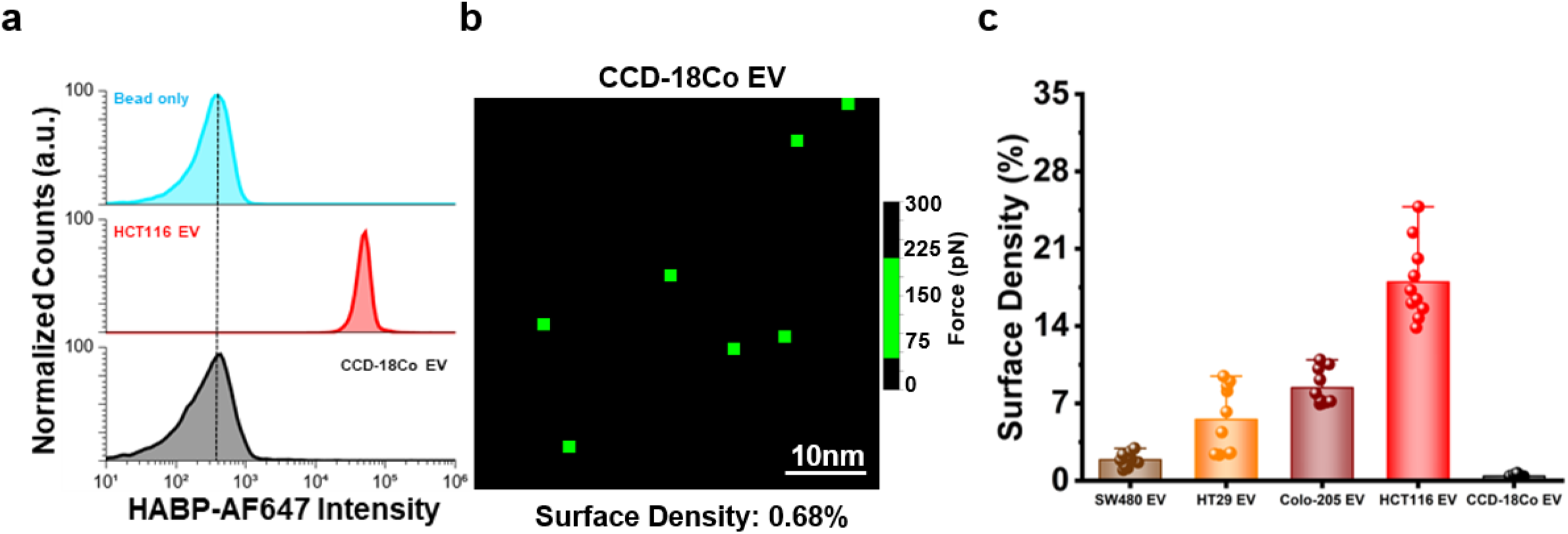
Control experiment. (a) Flow cytometry analysis quantifying surface-associated HA levels on CCD-18Co–derived EVs, shown in comparison with EVs from HCT116. (b) Representative force-volume adhesion map acquired on CCD-18Co–derived EV using HABP-functionalized AFM probe. (c) HA surface density on CCD-18Co–derived EVs compared with other EVs.

**Figure S10:**
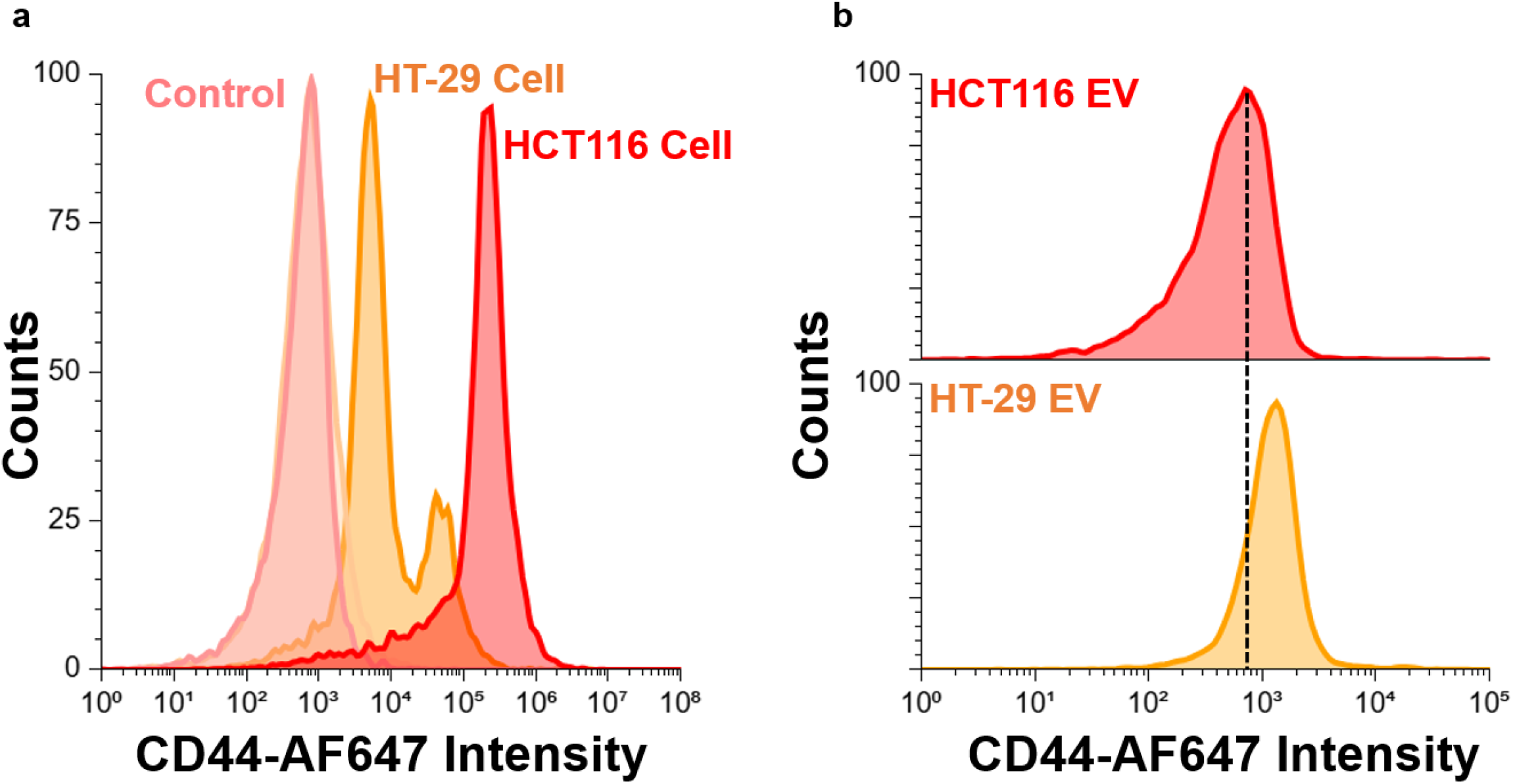
Flow cytometry–based quantification of CD44 expression in (a) stage B (HT29) and stage D (HCT116) cells and their (b)EVs.

**Figure S11:**
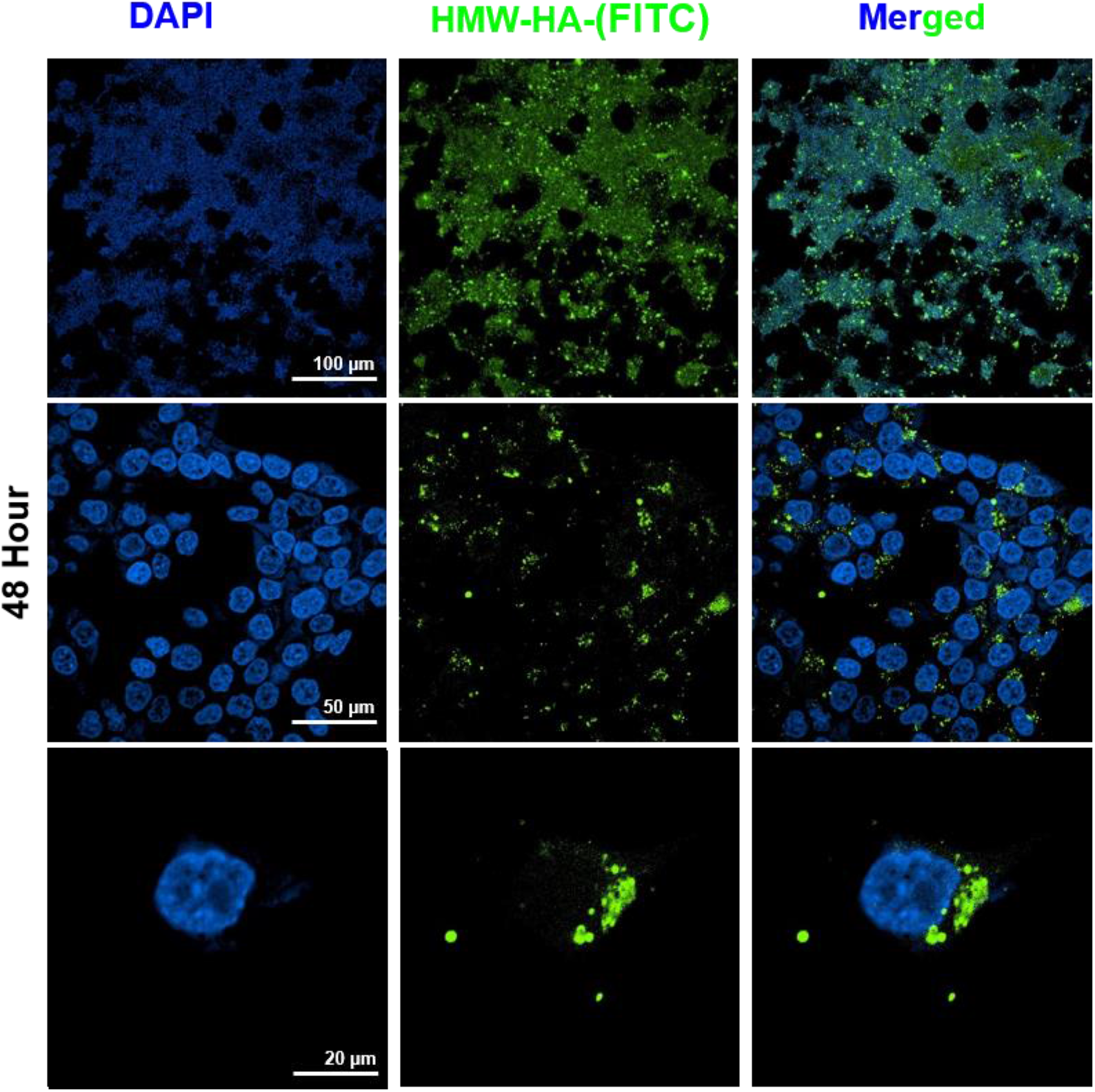
Visualization of exogenously added HMW-HA on stage D HCT 116 cells. Representative confocal images showing surface-associated FITC-tagged HMW-HA in green after 48 h of incubation. Images are displayed at progressively increasing magnification from top to bottom to highlight the spatial distribution and surface localization of HMW-HA.

**Figure S12:**
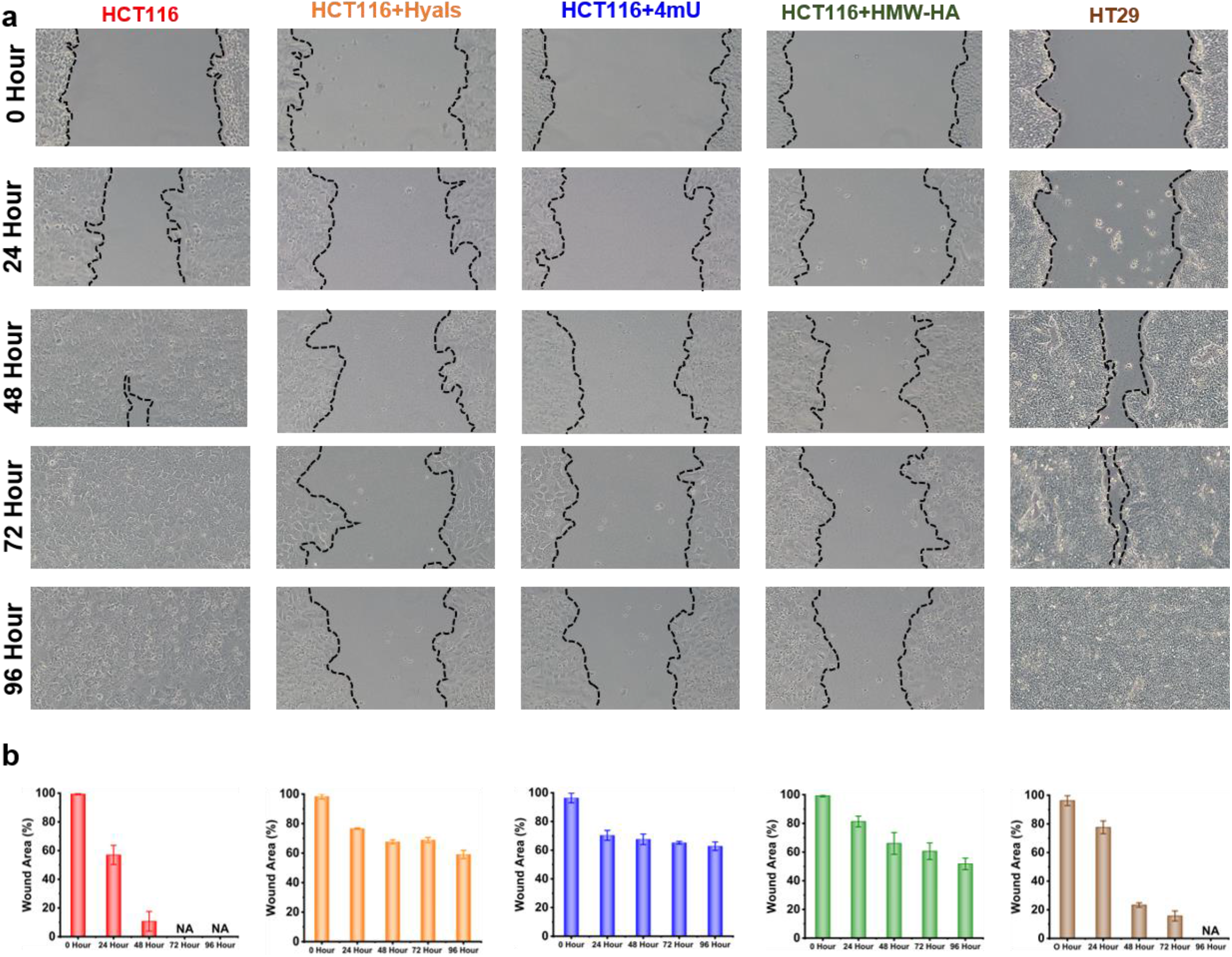
Effect of HMW-HA supplement, HA degradation, and HA synthesis inhibition on the migratory capacity of cells. (a) Representative phase-contrast images showing scratch closure in HCT116 cells and treated cells with HMW-HA hyaluronidase, or 4-MU at 0-96 h , stage B HT29 cells included as a reference (b) Quantitative analysis of scratch healing rates corresponding to the images.

**Figure S13:**
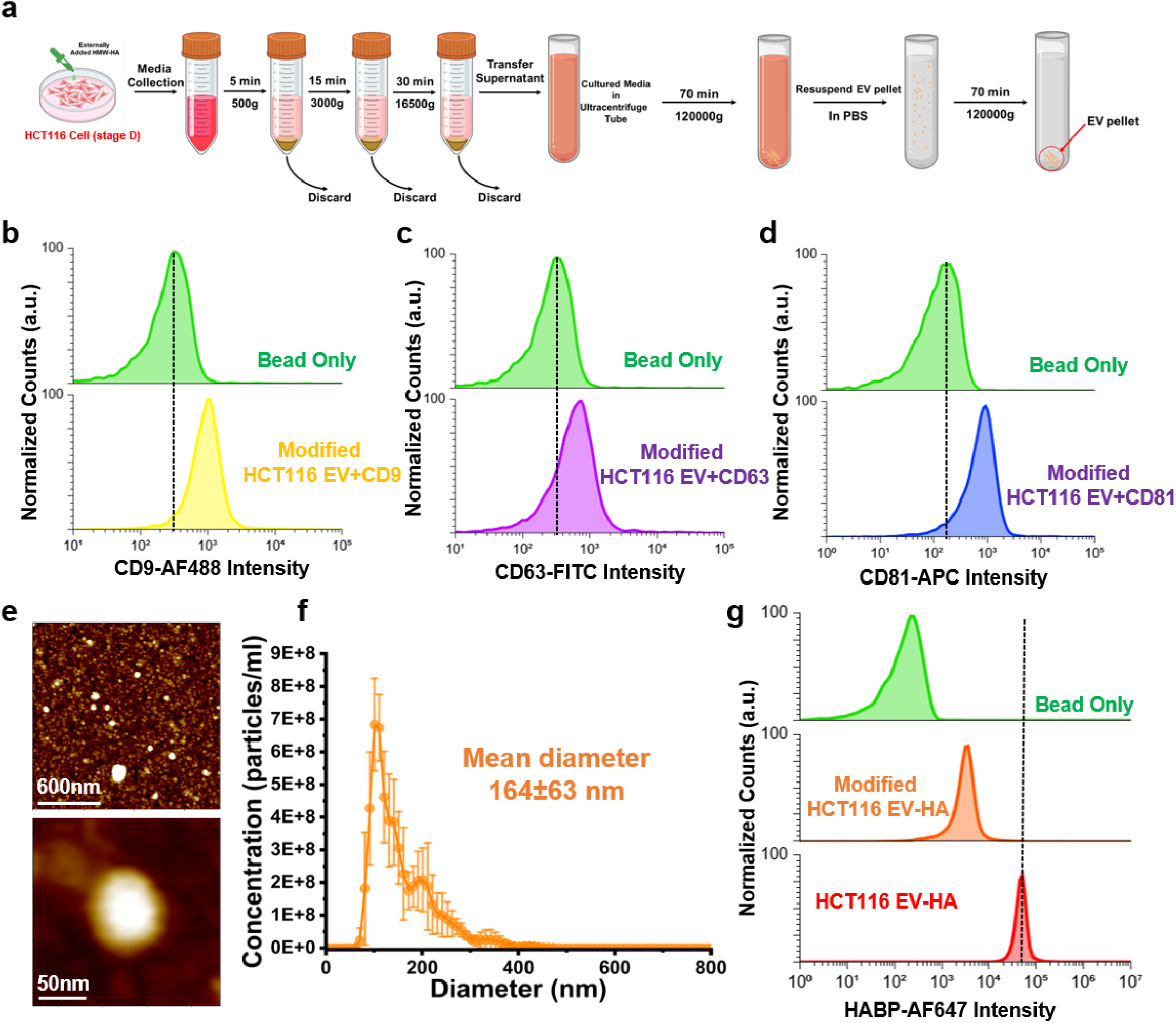
EVs derived from HMW-HA–modified stage D HCT116 CRC cells. (a) Schematic representation of the differential ultracentrifugation protocol used to isolate EVs from conditioned culture media of HMW-HA–treated stage D HCT116 cells (b–d) Flow cytometry of EV tetraspanin markers CD9, CD63, and CD81, respectively, (e) Representative AFM images revealing the spherical morphology of EVs isolated from HMW-HA–modified HCT 116 cells. (f) NTA showing the size distribution and particle concentration of isolated EVs, (g) Flow cytometry of surface-associated HA on EVs derived from HMW-HA–modified HCT 116 cells.

**Figure S14:**
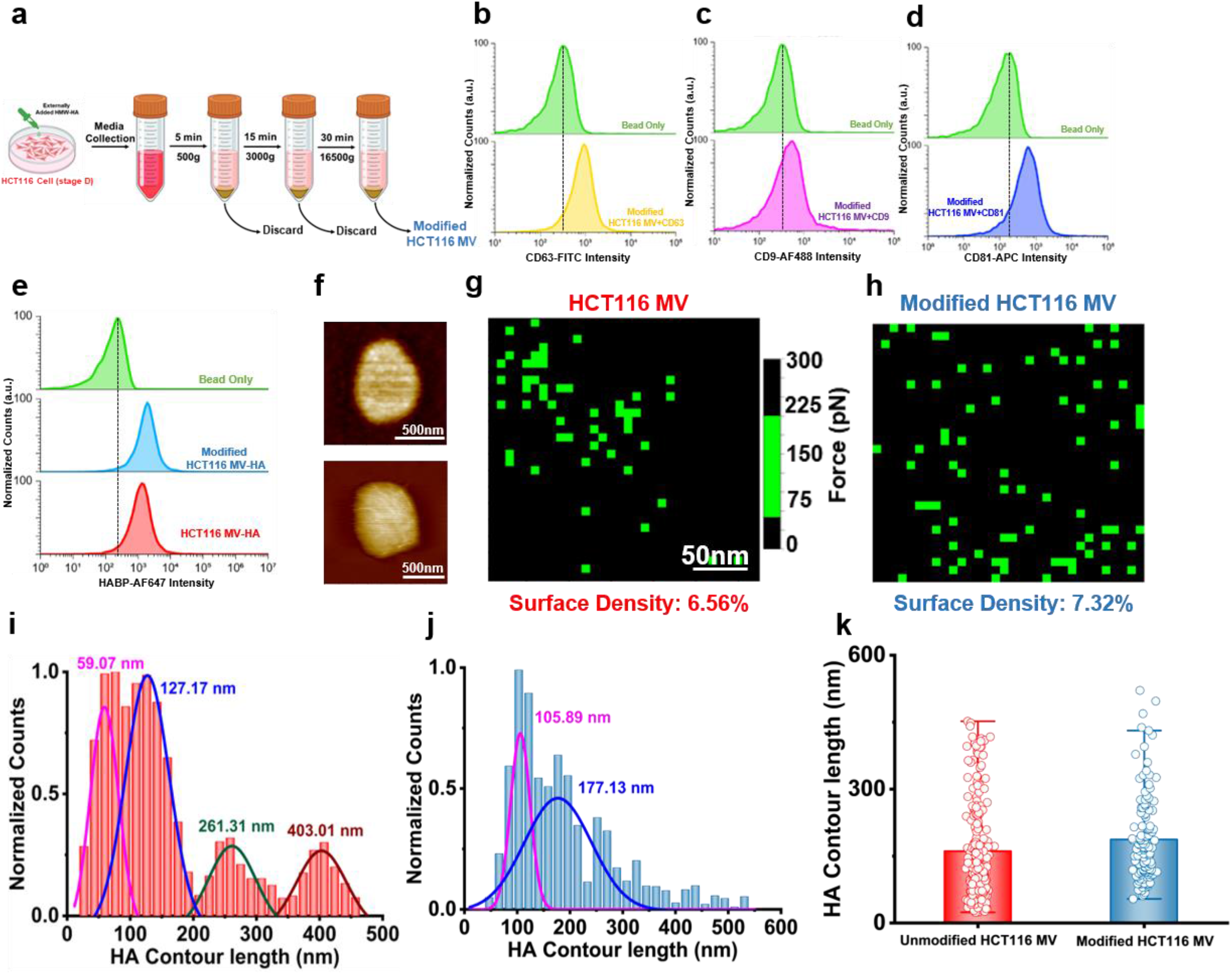
HA surface density, mechanical properties, and chain length on MVs derived from HMW-HA–modified HCT 116 cells. (**a**) Schematic overview of the stage HCT116 surface modification with HMW-HA and the differential ultracentrifugation workflow employed for isolation of MVs from treated cells. (**b–d**) Flow cytometry analysis showing tetraspanin markers CD9, CD63, and CD81, respectively. (**e**) Flow cytometry quantification of surface-associated HA on MVs isolated from HMW-HA–HCT116 cells (**f**) Representative AFM images of near-spherical morphology of MVs. (g, h) Representative AFM force–volume adhesion maps acquired on individual MVs. (**i-j**) Probability distributions of HA contour lengths measured on MVs isolated from modified and unmodified HCT116 Cells. (**k**) Comparison of the overall HA contour lengths.

**Table S1:** Physical properties of cells and EVs.

| Parameter | HT29 Cells (Stage B) | SW480 Cells (Stage B) | SW620 Cells (Stage C) | Colo-205 Cells (Stage D) | HCT116 Cells (Stage D) | Modified HCT116 Cells | HT29 EVs (Stage B) | SW480 EVs (Stage B) | SW620 EVs (Stage C) | Colo-205 EVs (Stage D) | HCT116 EVs (Stage D) | Modified HCT116 EVs | HT29 MVs (Stage B) | HCT116 MVs (Stage D) | Modified HCT116 MVs |
| --- | --- | --- | --- | --- | --- | --- | --- | --- | --- | --- | --- | --- | --- | --- | --- |
| Height | 2.99 ± 0.5µm | - | - | - | 3.69 ± 0.6µm | - | 15.2 ± 1 nm | 16.9 ± 3 nm | 23.7 ± 4 nm | 17.5 ± 3 nm | 20.7 ± 5 nm | 15.7 ± 2.3 nm | 107.3 ± 40 nm | 115.4 ± 20 nm | 47.3 ± 9 nm |
| Diameter (nm) | - | - | - | - | - | - | 58.5 ± 12 nm | 63 ± 5 nm | 95 ± 17 nm | 71 ± 11 nm | 86.5 ± 15 nm | 82.8 ± 3 nm | 314 ± 49 nm | 437.5 ± 94 nm | 357.1 ± 52 nm |
| Surface Area | 250 ± 61 µm² | - | - | - | 403 ± 132 µm² | 546 ± 139 µm² | 2062 ± 620 nm² | 3159 ± 547 nm² | 10502 ± 3162 nm² | 4046 ± 1359 nm² | 6065 ± 2124 nm² | 4649 ± 1985 nm² | 80700 ± 26054 nm² | 171077 ± 72231 nm² | 102182 ± 30040 nm² |
| Roughness (nm) | 55 ± 14 | - | - | - | 48 ± 12 | - | 0.64 ± 0.28 | 0.72 ± 0.19 | 0.92 ± 0.18 | 1.42 ± 0.44 | 2.44 ± 0.69 | 0.78 ± 0.12 | 1.08 ± 0.26 | 1.82 ± 0.58 | 1.06 ± 0.22 |
| Pericellular Coat Thickness (µm) | 1.01 ± 0.58 | - | - | - | 1.63 ± 0.66 | 2.56 ± 0.79 | - | - | - | - | - | - | - | - | - |
| HA Surface Density (%) | 3.17 ± 0.4 | 0.56 ± 0.1 | 3.9 ± 0.7 | 6.7 ± 2.7 | 10.6 ± 1.4 | - | 5.5 ± 2.8 | 3.7 ± 0.8 | - | 9.3 ± 4.8 | 17.9 ± 3.3 | 13.1 ± 3 | 4.5 ± 0.8 | 6.1 ± 1 | 5.8 ± 1 |
| Total HA per Unit Surface | 130146 ± 31661 | - | - | - | 700306 ± 229166 | - | 46 ± 13 | 44 ± 8 | - | 141 ± 47 | 445 ± 155 | 249 ± 106 | 58.1 ± 18 | 171 ± 72 | 81.8 ± 24 |
| LMW-HA (<200 kDa; <527 nm) (%) | 3.79% | 14.15% | 50.15% | 100% | 100% | 76.70% | 89% | 95% | - | 100% | 100% | 84.26% | 99.26% | 100% | 100% |
| HMW-HA (>200 kDa; >527 nm) (%) | 96.2% | 85.85% | 49.85% | 0% | 0% | 31.90% | 11% | 5% | - | 0% | 0% | 15.73% | 0.74% | 0% | 0% |
| Young's modulus | 1.61 ± 0.3 kPa | 1.43 ± 0.2 kPa | - | 0.59 ± 0.1 kPa | 0.72 ± 0.1 kPa | - | 2.6 ± 0.3 MPa | 2.4 ± 0.3 MPa | 1.16 ± 0.2 MPa | 1.6 ± 0.2 MPa | 1.9 ± 0.3 MPa | 3.5 ± 0.5 MPa | - | - | - |
| Viscosity (Pa-s) | 691 ± 18 | 698 ± 41 | - | 235 ± 8 | 334 ± 12 | - | - | - | - | - | - | - | - | - | - |
| Concentration (particles/ml) | - | - | - | - | - | - | 2x10<sup>10</sup> ± 7x10<sup>8</sup> | 4x10<sup>10</sup> ± 2x10<sup>9</sup> | 1.7 x 10<sup>11</sup> ± 6.8 x 10<sup>9</sup> | 6x10<sup>10</sup> ± 3x10<sup>9</sup> | 6.5x10<sup>10</sup> ± 3x10<sup>9</sup> | 5.6x10<sup>10</sup> ± 8x10<sup>8</sup> | - | - | - |
| NTA Diameter (nm) | - | - | - | - | - | - | 146 ± 65 | 139 ± 59 | 150 ± 55 | 128 ± 51 | 186 ± 68 | 154 ± 63 | - | - | - |

**Table S2:** Summary of the number of biological replicates, number of measured cells/EVs/MVs (n), and the total number of force-distance curves analysed.

| **System** | **Number of Biological Replicates** | **n** | **Force Curve Analyzed** |
| --- | --- | --- | --- |
| **HT-29 Cell** | **3** | **10** | **10216** |
| **SW480 Cell** | **3** | **10** | **9192** |
| **SW620 Cell** | **3** | **10** | **8192** |
| **Colo-205 Cell** | **3** | **10** | **11240** |
| **HCT116 Cell** | **3** | **10** | **10216** |
| **HT-29 EV** | **3** | **10** | **16444** |
| **SW480 EV** | **3** | **10** | **16363** |
| **SW620 EV** | **3** | **10** | **4596** |
| **Colo-205 EV** | **3** | **10** | **12448** |
| **HCT116 EV** | **3** | **10** | **17608** |
| **Mod-HCT116 EVs** | **3** | **10** | **16505** |
| **HT-29 MV** | **3** | **10** | **7168** |
| **HCT116 MV** | **3** | **10** | **8192** |
| **Mod-HCT116 MV** | **3** | **10** | **9126** |

**Table S3:** Model and simulation parameters (reduced units, *m* = *σ* = *k_B_T* = 1). Pair-interaction strengths are labelled by the interacting species: p = polymer, m = membrane, v = vesicle.

| Quantity | Symbol | Value |
| --- | --- | --- |
| **Membrane potential** |  |  |
| Interaction strength | $\epsilon$ | $4.5 k_B T$ |
| Preferred separation | $r_{\min}$ | $2^{1/6} \sigma \approx 1.122 \sigma$ |
| Cutoff radius | $r_c$ | $2.6 \sigma$ |
| Fluidity exponent | $\zeta$ | 4.0 |
| Bending-rigidity parameter | $\mu$ | 3.0 |
| Spontaneous-curvature angle | $\Theta_0$ | 0.0 |
| **Grafted polymers** |  |  |
| Backbone bond stiffness | $K_{\text{bond}}^{\text{bb}}$ | 100 |
| Graft bond stiffness | $K_{\text{bond}}^{\text{graft}}$ | 200 |
| Bond rest length | $r_0$ | $1.0 \sigma$ |
| Backbone bending stiffness | $K_{\text{angle}}^{\text{bb}}$ | 8.0 |
| Graft bending stiffness | $K_{\text{angle}}^{\text{graft}}$ | 12.0 |
| **Non-bonded Lennard-Jones interactions** |  |  |
| Polymer-membrane (adhesive) | $\epsilon_{\text{pm}}, r_c$ | $2.2 k_B T, 2.5 \sigma$ |
| Polymer-polymer (WCA) | $\epsilon_{\text{pp}}, r_c$ | $1.0 k_B T, 2^{1.6} \sigma$ |
| Membrane-vesicle (WCA) | $\epsilon_{\text{mv}}, r_c$ | $1.0 k_B T, 2^{1.6} \sigma$ |
| Vesicle-polymer (WCA) | $\epsilon_{\text{vp}}, r_c$ | $1.0 k_B T, 2^{1.6} \sigma$ |
| **Geometry and run control** |  |  |
| Vesicle radius | $R$ | $10 \sigma$ |
| Polymer length | $L$ | $10\text{-}40 \sigma$ |
| Grafting surface density | $\Phi_s$ | 5-20% |
| Production length | - | $2\text{-}4 \times 10^7$ steps |

